# Molecular determinants of avoidance and inhibition of *Pseudomonas aeruginosa* MexB efflux pump

**DOI:** 10.1101/2023.06.01.543207

**Authors:** Silvia Gervasoni, Jitender Mehla, Charles Bergen, Inga V. Leus, Enrico Margiotta, Giuliano Malloci, Andrea Bosin, Attilio V. Vargiu, Olga Lomovskaya, Valentin V. Rybenkov, Paolo Ruggerone, Helen I. Zgurskaya

**Author notes:** Broad Institute of MIT and Harvard, Center for the Development of Therapeutics, 415 Main Street, Cambridge, MA 02142, United States.

## Abstract

Transporters of the Resistance-Nodulation-cell Division (RND) superfamily of proteins are the dominant multidrug efflux power of Gram-negative bacteria. The major RND efflux pump of *Pseudomonas aeruginosa* is MexAB-OprM, in which the inner membrane transporter MexB is responsible for recognition and binding of compounds. The high importance of this pump in clinical antibiotic resistance made it a subject of intense investigations and a promising target for the discovery of efflux pump inhibitors. This study is focused on a series of peptidomimetic compounds developed as effective inhibitors of MexAB-OprM. Previous analyses of antibacterial and biochemical activities showed that these compounds vary broadly in their efficiency as inhibitors or substrates of MexAB and can be categorized into different functional classes. Here, we performed multi-copy molecular dynamics simulations, machine learning analyses and site-directed mutagenesis of MexB to investigate interactions of MexB with representatives of the various classes. The analysis of both direct and water-mediated protein-ligand interactions revealed characteristic patterns for each class, highlighting significant differences between them. We found that efflux avoiders poorly interact with the access binding site of MexB, and inhibition engages amino acid residues that are not directly involved in binding and transport of substrates. In agreement, machine learning models selected different residues predictive of MexB substrates and inhibitors. The differences in interactions were further validated by site-directed mutagenesis. We conclude that the substrate translocation and inhibition pathways of MexB split at the interface (between the main putative binding sites) and at the deep binding pocket, and that interactions outside of the hydrophobic patch contribute to the inhibition of MexB. This molecular-level information could help in the rational design of new inhibitors and antibiotics less susceptible to the efflux mechanism.

**Importance:** Multidrug transporters recognize and expel from cells a broad range of ligands including their own inhibitors. The difference between the substrate translocation and inhibition routes remains unclear. In this study, machine learning, computational and experimental approaches were used to understand dynamics of MexB interactions with its ligands. Our results show that some ligands engage a certain combination of polar and charged residues in MexB binding sites to be effectively expelled into the exit funnel, whereas others engage aromatic and hydrophobic residues that slow down or hinder the next step in the transporter cycle. These findings suggest that all MexB ligands fit into this substrate-inhibitor spectrum depending on their physico-chemical structures and properties.

## Introduction

*Pseudomonas aeruginosa* is an opportunistic Gram-negative pathogen responsible for infections associated with high morbidity and mortality rates (1)(2)(3). The lack of effective antimicrobials against this bacterium arises from different resistance mechanisms, among which the action of efflux pumps represents one of the major contributors (4)(5)(6)(7). The tripartite efflux system MexAB-OprM of *P. aeruginosa* plays the leading role in translocating a plethora of different antimicrobial compounds outside the cell (8)(9)(10). The homotrimeric protein MexB is a Resistance Nodulation cell Division (RND) transporter responsible for the binding and extrusion of ligands (11)(12) through the so-called functional rotation mechanism (5)(13)(14). In this mechanism, the protomers can assume cyclically three different and subsequent states, namely Loose (L), Tight (T) and Open (O). The substrate-transport process is energetically driven by the electrochemical potential of protons, i.e., the proton motive force (PMF). The pump expels compounds with very different physico-chemical properties, a feature known as poly-specificity (10)(15), which is shared by many RND transporters from Gram-negative species including AcrB from *Escherichia coli* (16) or AdeB from *Acinetobacter baumannii* (17), both homologous to MexB. Based on the available structural data of these transporters, two main binding pockets named Access Pocket (AP_L_) and Distal Pocket (DP_T_) have been identified in the L and T states, respectively (11)(18)(19)(20). These two pockets are separated by the interface, which includes the switch (gate) loop (**Figure S1**). The composition and the size of the loop are thought to contribute to substrate specificity of RND transporters (21). Significant efforts have been made to enhance the efficacy of antibacterial strategies against *P. aeruginosa* and other Gram-negative bacteria (9)(23)(24)(25), but the insufficient progress indicates that innovative approaches are urgently needed.

The most desirable feature of new antimicrobial compounds is the ability to avoid efflux. Therefore, to design effective drugs, it is of great importance to identify good and poor efflux substrates (24)(23)(26)(27), and the molecular determinants guiding avoidance (28). In addition, a promising strategy is represented by the development of molecules able to inhibit RND transporters, leading to an effective accumulation of antibiotics inside the cell (29). Several efflux pump inhibitors (EPIs) have been identified (30) such as ABI-PP (18) and the peptidomimetics PaβN (30) for MexB, as well as the MBX series (31), 4-substituted 2-naphthamide derivatives (32) and natural compounds (33)(34) for AcrB. Peptidomimetics raised great interest thanks to their promising broad-spectrum activity (28). Two mechanisms of inhibition have been reported: allosteric and competitive. In the former, as seen for AcrB, the inhibitor binds at the transmembrane domain of the L protomer, which would prevent either the transition from L to T states or the use of PMF for substrate efflux (35). Competitive inhibitors bind at the same sites of substrates, hindering their capture and subsequent translocation (36)(37). According to experimental and structural data, some EPIs bind within a peculiar region inside the DP_T_, known as a hydrophobic trap (5)(18), preventing the binding of substrates. On the other hand, most of the identified EPIs are also substrates of efflux pumps.

Although the use of EPIs has shown several advantages(38), none of the reported EPIs has been approved for clinical use so far, mainly because of the low *in vivo* efficacy, poor pharmacokinetic properties and/or toxicity problems. Further understanding of molecular mechanisms associated with MexB recognition, inhibition and possibly avoidance, could facilitate the development of EPIs suitable for clinical applications.

In a previous work (28), ∼260 peptidomimetics developed by Rempex Pharmaceuticals (Rempex compounds) were characterized through growth-dependent and -independent structure-activity-relationship (SAR) analyses and classified into four functional groups: substrates (SUBs, 136 molecules), inhibitors (EPIs, 24 molecules), mixed substrates-inhibitors (EPI-Ss, 31 molecules), and avoiders (AVDs, 66 molecules). This series of compounds are the most advanced among various EPIs and contain compounds with good pre-clinical profiles (39). The AVDs were defined as molecules that are not translocated by MexB and, unlike the inhibitors, they do not even bind the transporter. The mixed EPI-Ss can inhibit the efflux of substrates, but at the same time, they can be transported by MexB as well. Physico-chemical properties of compounds and their interactions with MexB using molecular docking were computed. These descriptors were used to generate Machine-Learning (ML) models that are predictive of the propensity of compounds to avoid or inhibit efflux by MexB. Several descriptors of interactions with MexB were among the top predictors of whether a compound would be recognized by MexB as a ligand, or it will avoid the transporter. In particular, the strong affinity to AP_L_ and the higher number of contacts with L674 and P668 residues in this pocket as well as with T130, F136 and S276 in DP_T_ correlated positively with the activities of EPIs. However, the generality of the model predictions was not always clear, prompting for a more mechanistically focused analysis.

In this study, to gain a mechanistic understanding of differences between substrates and EPIs in their interactions with MexB, we performed improved ML analysis of MexB residues located in ligand binding pockets, multi-copy molecular dynamics (MD) simulations of four Rempex compounds that are representative of the four functional classes, and site-directed mutagenesis of specific MexB residues implicated in interactions with ligands. The ML analysis and the MD simulations showed that MexB interactions predictive of substrates span multiple binding sites in AP, DP, and the interface between the two pockets, whereas MexB residues/regions important for inhibition are localized specifically to DP. However, interactions with the hydrophobic trap alone are not sufficient for inhibition and amino acid residues from other regions of DP contribute to inhibition. In agreement with previous studies (5)(40)(41), we found that solvent water molecules play an important role in binding and inhibition. Site-directed mutagenesis of the identified MexB residues confirmed their functional importance and specificity in inhibition and translocation. The identified MexB residues selective for substrates and inhibitors will improve predictions of these properties in structurally diverse compound libraries.

## Results

### Machine Learning analysis of interaction counts to MexB residues located in ligand binding pockets

To quantify the interactions of Rempex compounds with MexB at a molecular level, we first carried out ensemble docking calculations as described in previous works (28)(42). An extended description of the docking results is reported in the **Supporting Information**. Briefly, we generated 600 poses per ligand in each of the two major putative binding pockets of MexB (i.e., AP_L_ and DP_T_), which were further subdivided into the Outer and Inner AP, the Interface (IF), which includes the switch loop (residues 613-623), and the DP Groove and DP Cave regions (**Figure 1A**). Residues lining the five different regions of MexB (Outer AP, Inner AP, Interface, DP Groove and DP Cave) are listed in **Table S1**.

**Figure 1.**
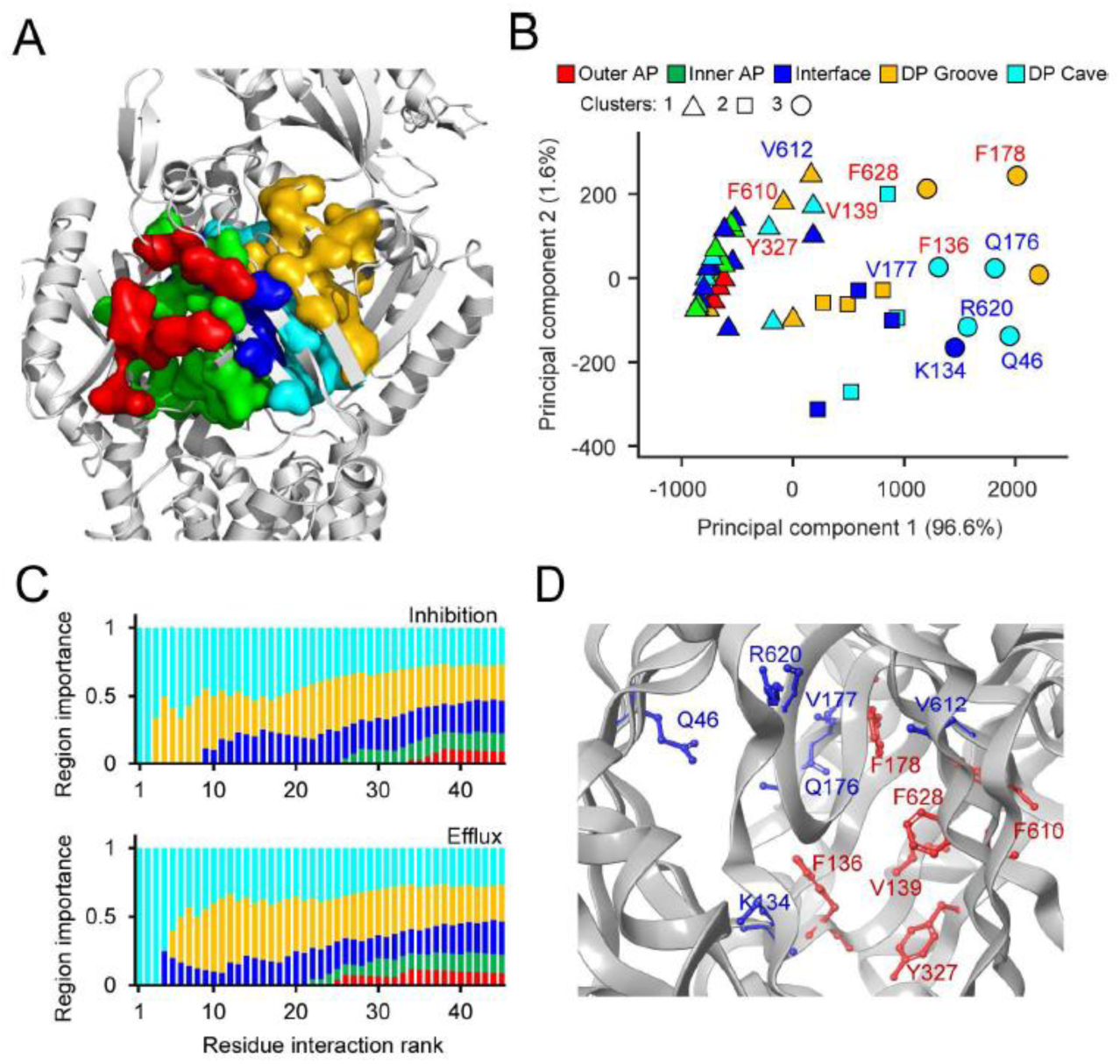
Machine Learning analysis of MexB interaction with substrates and inhibitors. **A.** Architecture of the substrate binding site of MexB with its five main areas highlighted as following: Outer AP-red, Inner AP – green, Interface – blue, DP Groove – gold; DP Cave – cyan. **B.** Principal component analysis of residue interaction counts. Symbols denote the three clusters predicted by hierarchical clustering, colours mark location of the residues in the binding sites of MexB. The top six residues predicted by the random forest analysis are labelled in red for inhibition and blue for efflux efficiency. **C.** Representation of the relative importance of the five main MexB regions among top residue interaction counts that were ranked using a random forest analysis and then grouped according to their location within MexB.

Hierarchical clustering analysis of all 206 Rempex compounds and their contacts with MexB showed that the MexB residues form three clusters (**Figure 1B**). The cluster 1 comprised all the residues of the Outer and Inner AP and a few residues from the IF and the DP Cave, whereas the clusters 2 and 3 were broadly dispersed and included residues from the DP Groove, the DP Cave, and the IF. Thus, clusters are not tightly associated with specific regions of MexB structure, although there is a clear separation between the interaction in the AP and DP. The Principal Component decomposition (43) showed that the first and second principal components (PComp1 and PComp2) account for 96.6% and 1.6% of the total variance, respectively (**Figure 1B**). Notably, most of the variance in the dataset was explained by PComp1, which included almost equal (varied within a factor of 2.4) contributions from all compounds. Thus, most of the variation among the compounds is associated with their overall propensity for interaction with the residues of MexB. Interestingly, the PComp1 clearly separates cluster 1 (hence, the AP) from the rest of the MexB interactions.

We next applied a further ML analysis to identify MexB residues predictive of substrates and inhibitors. To generate a model for MexB substrates, we used the previously reported efflux ratios of compound concentrations inhibiting 50% of bacterial growth (IC_50_) measured in efflux-proficient *P. aeruginosa* PAO1(Pore) and efflux-deficient PΔ6(Pore) cells (28). The MexB substrates were defined as compounds with efflux ratios ≥ 4. The EPI activities were determined using the inhibition of efflux of a fluorescent probe Hoechst 33342 (Hoechst) in non-growing bacterial cells. The ratio of Hoechst accumulation rates at 16 μM and 0 μM concentrations of a compound was used as a proxy for its efficiency as an inhibitor. The two ratios were modelled as a random forest ensemble of regression trees using the docking-generated residue interaction counts as descriptors. The generated models were used as a basis for ranking the descriptors according to their importance for the classification of compounds.

The highest ranked amino acid residues of MexB lie in the DP Cave and DP Groove sections followed by the IF region (**Figure 1C**). Notably, the overall ranking profiles were very similar for efflux substrates and inhibitors. This lends further support to the accepted notion that the site for MexB inhibition is orthosteric with its normal efflux site. However, different residues were found at the top of the lists of the two classes of molecules. The top six residues most important for the inhibition were all aromatic or hydrophobic (F136, V139, F178, F610, Y327, and F628) whereas the efficiency of efflux was defined by interactions with primarily polar residues (Q46, K134, Q176, V177, V612 and R620) (**Figure 1B**). Thus, both DP Cave and DP Groove appear important to discriminate between different classes of ligands, i.e. substrates vs. avoiders and inhibitors vs. non-inhibitors. However, the propensity to inhibition is largely defined by interactions with aromatic and hydrophobic residues.

### Molecular docking suggests common binding modes for efflux substrates, inhibitors, and avoiders

We next performed a detailed structural analysis of the top docking poses found in each binding pocket for representative compounds of the four Rempex classes: SUB58, EPI18, mixed EPI-S32, and AVD108 (**Figure 2, Table S2**). SUB58 and EPI-S32 have only weak antibacterial activities with MICs values ≥100 μM and 25 μM against PA2859 (Pore) producing MexB WT and carrying an empty vector, respectively. The MIC values of AVD108 and EPI18 were 6. 25 μM and 12.5 μM, respectively, against both MexB WT and null cells (**Table S2**).

**FIGURE 2:**
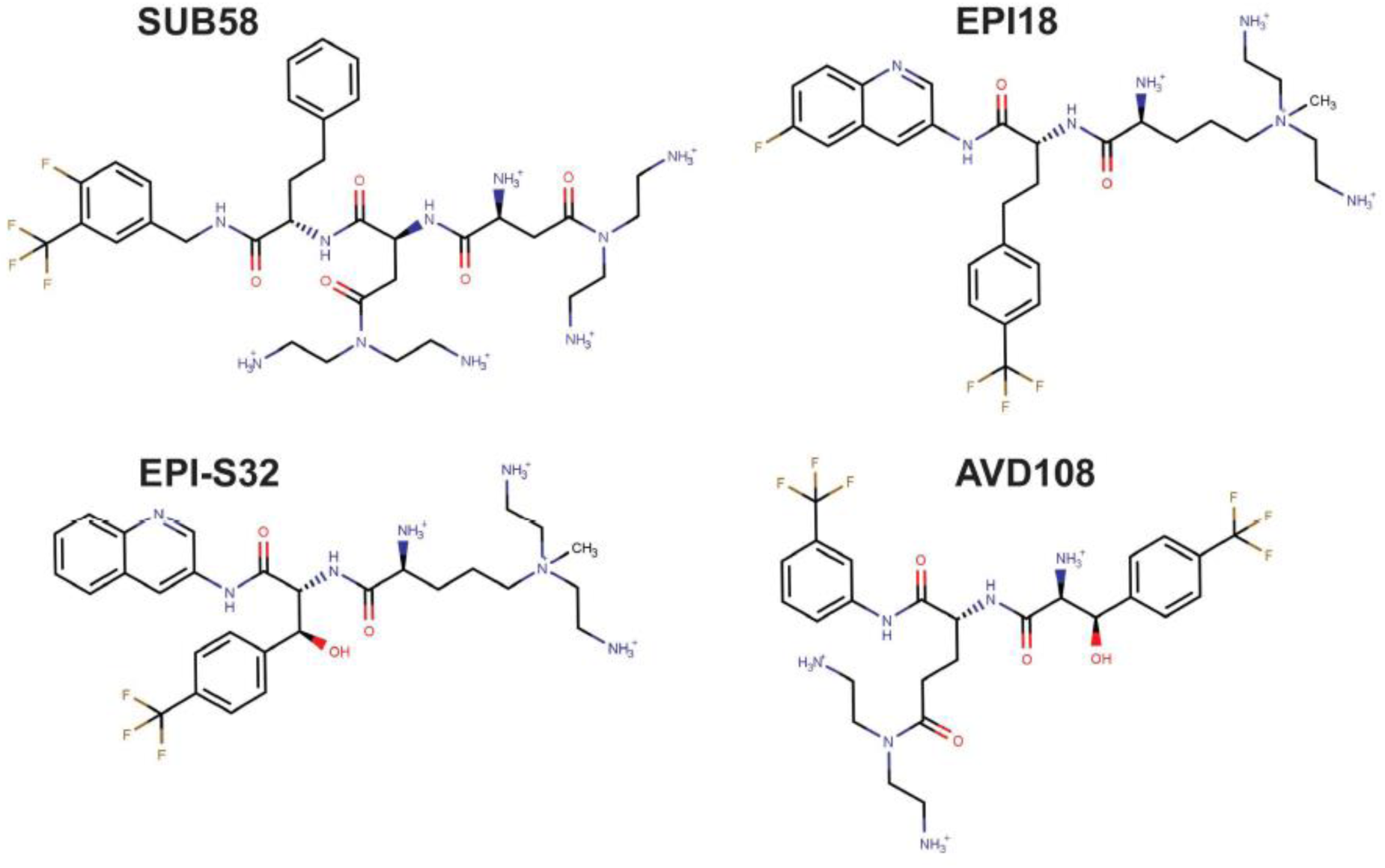
Chemical structures of the representative compounds of the four Rempex classes: SUB58, EPI18, EPI-S32, and AVD108.

Overall, the analysis of the docking poses reveals similar binding modes for the representative Rempex compounds in the putative binding sites of MexB. The differences between substrates, inhibitors and avoiders are subtle and appear at the level of specific residues involved in the binding. In particular: docking poses at the AP_L_ (**Figure S2** and **Figure 3**) reveal a common binding mode, i.e., the hydrophobic rings of compounds point towards the Inner AP and IF, containing apolar residues such as L564, P669, V671, L674 and L861; the terminal amines are associated with the Outer AP. This region is responsible for most of the interactions between MexB and the Rempex compounds, although the residues involved vary to some extent. Docking poses at the DP_T_ (**Figure S3** and **Figure 4**) share the same orientation, with the polar tails facing the IF, and the aromatic rings directed toward the DP Cave. All ligands participate also in polar interactions with different residues through their positively charged amines, while EPI18 is involved in π-π stacking interactions with phenylalanine residues of the DP Groove.

**FIGURE 3.**
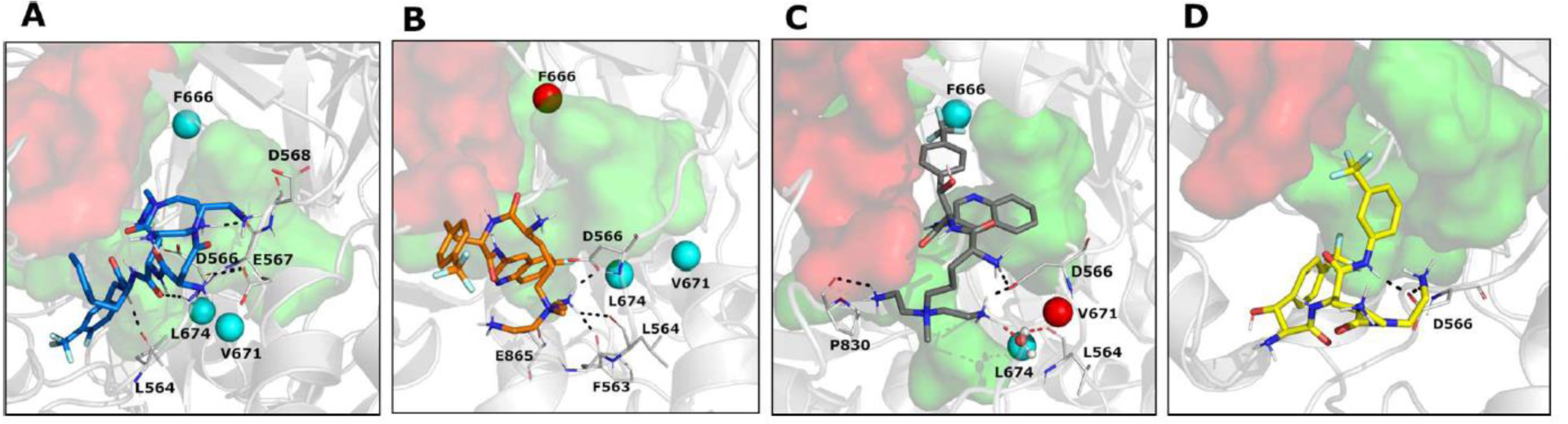
MD cluster representatives at the AP_L_. **A.** SUB58 is colored in blue (cluster population = 30%), **B.** EPI18 is in orange (cluster population = 29%), **C.** EPI-S32 is in grey (cluster population = 33%), and **D.** AVD108 is in yellow (cluster population = 41%). Protein-ligand interactions are highlighted as dotted lines and colored in black for hydrogen bonds and salt bridges, and in red for water H-bridges. In panel **C**, water molecule is represented in balls and sticks. MexB amino acid residues replaced by site-directed mutagenesis are shown as colored spheres: cyan – no change in function, green – loss of function and red-gain of function. The Outer AP and the Inner AP are highlighted in red and green surfaces, respectively.

**FIGURE 4.**
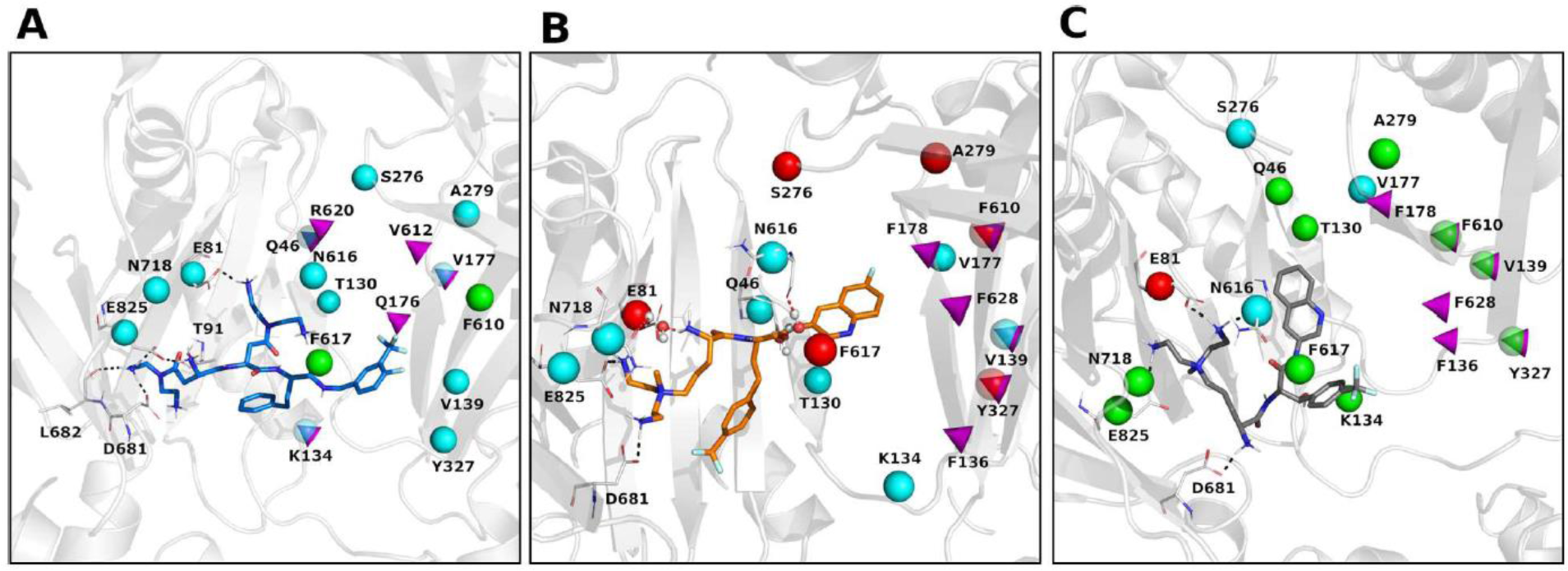
MD cluster representatives at the DP_T_. **A.** SUB58 is colored in blue (cluster population = 20%). **B.** EPI18 is in orange (cluster population = 41%). **C.** EPI-S32 is in grey (cluster population = 30%). Protein-ligand interactions are highlighted as dotted lines and colored in black for hydrogen bonds and salt bridges, and in red for water H-bridges. In panel **B**, water molecules are represented in balls and sticks. MexB amino acid residues replaced by site-directed mutagenesis are shown as colored spheres: cyan – no change in function, green – loss of function and red-gain of function. The DP Groove is encircled in blue, and the DP Cave is in red. Top predictors from ML analysis are highlighted as magenta cones.

Thus, the ML analyses of docking results show that interactions of Rempex compounds with the AP residues are distinct from those in the IF and DP. Compounds interact with the AP by facing their aromatic rings towards the inner part of the pocket, in a region delimited by hydrophobic residues (i.e., L564, P669, V671, L674 and L861), while the polycationic moieties face the periplasm and are exposed to the solvent.

### Substrates and inhibitors engage more MexB residues than efflux avoiders

To further assess docking findings and investigate the impact of solvent molecules in protein-ligand interactions, we performed all-atom MD simulations starting from the binding modes predicted at both AP_L_ and DP_T_. For each ligand-protein complex selected from docking (**Figures S2** and **S3**), we performed ten MD replicas of 100 ns (1 μs total sampling for each pose), by considering a phospholipid double membrane-embedded model of MexB in water solution (see **Methods** for further details). Our primary aim was to assess the binding modes predicted by docking for the representative compounds (i.e., SUB58, EPI18, EPI-S32 and AVD108) and infer any potential term of diversification between the four Rempex classes. Note that, consistently with its nature, the avoider AVD108 was simulated only in the AP_L_.

We first focused on the overall stability of the MD trajectories. In general, all compounds were stable in all simulations with some minor differences between them (**Table 1**, **Figure S4**).

**TABLE 1.**
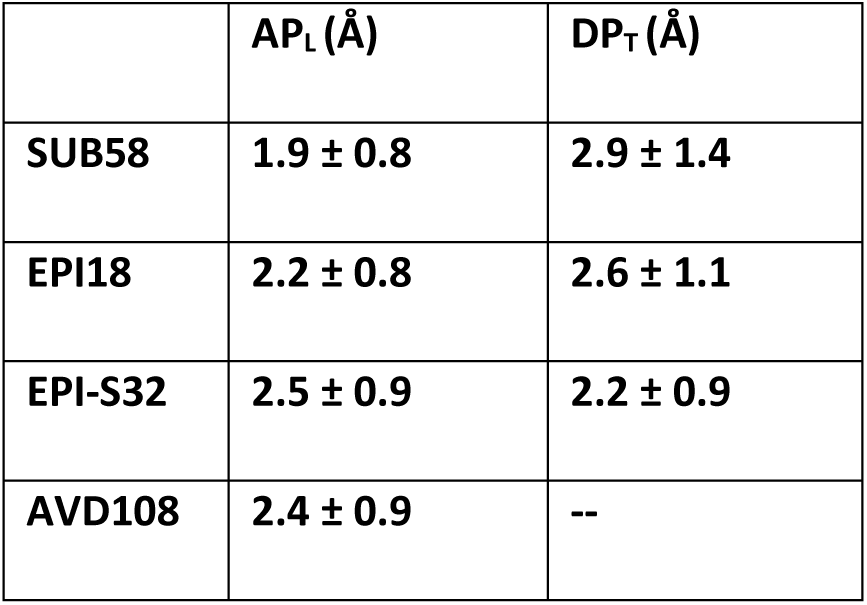
Average root-mean-square displacements (RMSD, expressed in Å) and associated standard deviations of the ten MD replicas performed for each representative Rempex compound at the AP_L_ and DP_T_.

In the AP_L_, the lowest and highest average RMSD values were found for the SUB58 and EPI-S32, respectively (1.9 ± 0.8 Å and 2.5 ± 0.9 Å). Conversely, SUB58 showed the highest RMSD values and EPI-S32 the lowest in the DP_T_ (2.9 ± 1.4 and 2.2 ± 0.9 Å, respectively). These higher RMSD values at the DP_T_ in the SUB58 simulations are consistent with the multisite-drug-oscillation (or diffusive binding) hypothesis (15)(44).

We then investigated the persistence of direct interactions of each compound with MexB residues, by monitoring the contacts along the MD trajectories (cutoff distance 3.0 Å, see **Methods**) (**Table S2**). Overall, we found that SUB58 interacted with the largest number of protein residues (especially at the Outer and Inner AP) as compared to the other compounds (**Figure 3**). Conversely, as expected, AVD108 showed the lowest values of persistence. Some of the residues appear to interact with all compounds (although to a different extent). For example, all compounds interact with D566 of the Outer AP with percentages 100%, 46%, 83%, and 47%, for SUB58, EPI18, EPI-S32, and AVD108, respectively (**Figure 3** and **Table S3**). Noteworthy, the interaction between D566 and SUB58 is constantly maintained for all the trajectories. The representative complexes extracted from MD simulations (**Figure 3**), reveal that interactions with the AP residues are mainly mediated by the amino-terminal tails and the amide moieties, which is consistent with previously reported findings (45).

Surprisingly, at the DP_T_ none of the compounds interacted with the Groove or Cave residues and all compounds primarily interacted with the residues located in the inner AP and the IF (**Figure 4**). All compounds established interactions with E81 (IF), D681 (Inner AP) and E825 (Inner AP), while unique interactions were found for SUB58 with T91 (IF) and L682 (Inner AP), and for EPI-S32 with N616 (IF) (**Figure 3**). Thus, MD simulations identify interactions important for individual compounds that are markers of high affinity binding and are involved in the translocation of all ligands. The rare contacts in the DP Groove that are important for distinguishing inhibitors from non-inhibitors can be captured by ML modelling fed with interaction descriptors computed on a larger series of compounds.

### Water-mediated hydrogen bonding contributes remarkably to the interaction with MexB

To assess differences in hydration profiles of compounds, we analyzed the water shells around the Rempex representatives both in complex with MexB, and alone in water solution as a reference term (see **Methods** and **Figure S5**). **Figure 5A** reports the average number of water molecules of the 1^st^ and 2^nd^ hydration shells in the MD trajectories performed at the AP_L_ and DP_T_, and for the sake of comparison the same numbers obtained for the compounds inserted in a water box (28). As expected, within the AP_L_ the compounds are surrounded by a higher number of water molecules as compared to the DP_T_. The only exception is represented by EPI18, which maintains similar hydration in both pockets. Interestingly, EPI18 shows the highest ratios as compared to the other compounds, and EPI-S32 showed a hydration profile like that of EPI18 (SUB58) at the AP_L_ (DP_T_), which is consistent with its mixed nature.

**FIGURE 5.**
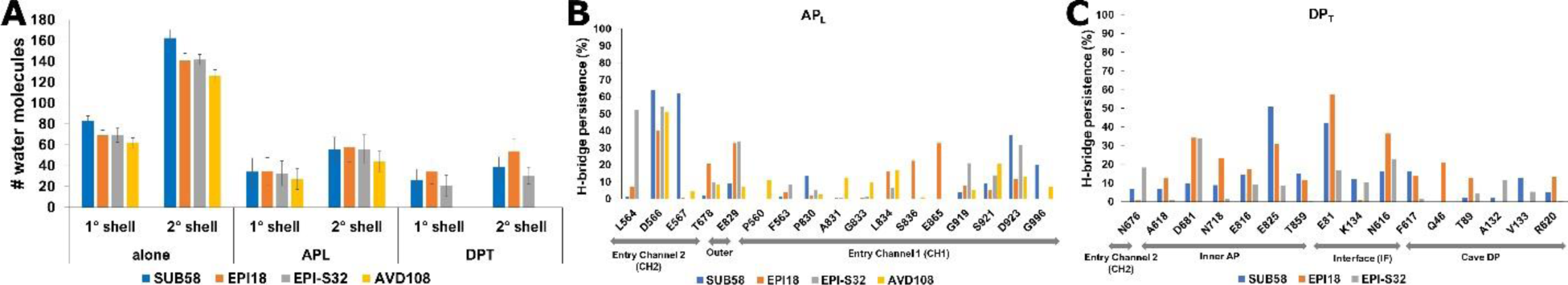
Compound hydration and water-mediated hydrogen bonding. **A.** Comparison between the average number of water molecules in the 1^st^ and 2^nd^ hydration shell registered in the MD trajectories of the Rempex compounds in complex with MexB at the AP_L_ and DP_T_, and alone in water solution. **B.** Mean values (%) of the persistence of hydrogen bond bridges mediated by water molecules during the MD trajectories, between Rempex compounds and MexB residues, at the AP_L_. **C.** Mean persistence of interaction (%) and corresponding standard deviation between MexB residues and the Rempex compounds, recorded in the DP_T_. We report only residues for which percentages greater than 12% were registered for at least one compound.

The above results suggest a prominent role of water molecules in the interaction between MexB and the representative Rempex compounds. Therefore, to further quantify the role of the solvent and pinpoint specific protein residues interacting indirectly with compounds, we computed the persistence of hydrogen bond bridges mediated by water molecules during the MD runs. H-bridges mediated by water molecules appear to contribute remarkably to the interaction with MexB (**Figure 5B-C**, **Table S4**). Furthermore, indirect water-mediated contacts involve residues lining the DP Cave of MexB, as seen for EPI18 and the Q46 and T89 residues. Overall, at the AP_L_, AVD108 is engaged in the lowest number of water bridges (17 H-bridges/ns, compared to 24, 22 and 27 of SUB58, EPI18 and EPI-S32, respectively). While at the DP_T_, EPI18 shows a marked tendency to interact with the water molecules (32 H-bridges/ns), compared to the other compounds (19 H-bridges/ns for both SUB58 and EPI-S32).

Thus, combining the results from the analysis of contacts (**Figures 3** and **4**) and water-mediated H-bridges (**Figure 5**), we found that the four classes of Rempex compounds engage MexB residues from different regions both directly and indirectly. For instance, at the AP_L_ the residue L564 of the Outer AP was directly interacting with SUB58 and EPI18 (**Figure 3**), but it was also indirectly interacting with EPI-S32 by a water H-bridge (**Figure 5B**). At the DP_T_, residues Q46 (DP Cave), N616 (IF) and N718 (Inner AP) are found to interact indirectly only with EPI18 (**Figure 5C**).

### Mutations in key MexB residues differentially affect substrate recognition

To gain insights into functional interactions of SUB58, EPI18, and EPI-S32 within the known binding pockets of MexB, and to further define properties that distinguish the translocation and inhibition pathways, we performed a site-directed mutational analysis. We considered a subset of key residues selected from the results of ML analysis and MD simulations described above (**Table 1** and **Figures 1, 3-5**), as well as residues previously implicated by structural studies in the substrate- and EPI recognition by MexB and its close homolog AcrB (**Figure 6**)(5)(46)(47). Among the selected residues, F666, L674, N718 and E825 are in the Inner AP, E81, K134, N616 and F617 are from the IF with the last two in the switch loop, Q46, T130, V139, V177 and Y327 are from the DP Cave and finally, S276, A279 and F610 are from the DP Groove (**Figure 1A**). Ten of the selected residues were replaced with cysteine (MexB-Cys), six with alanine and A279 was substituted with valine.

**Figure 6.**
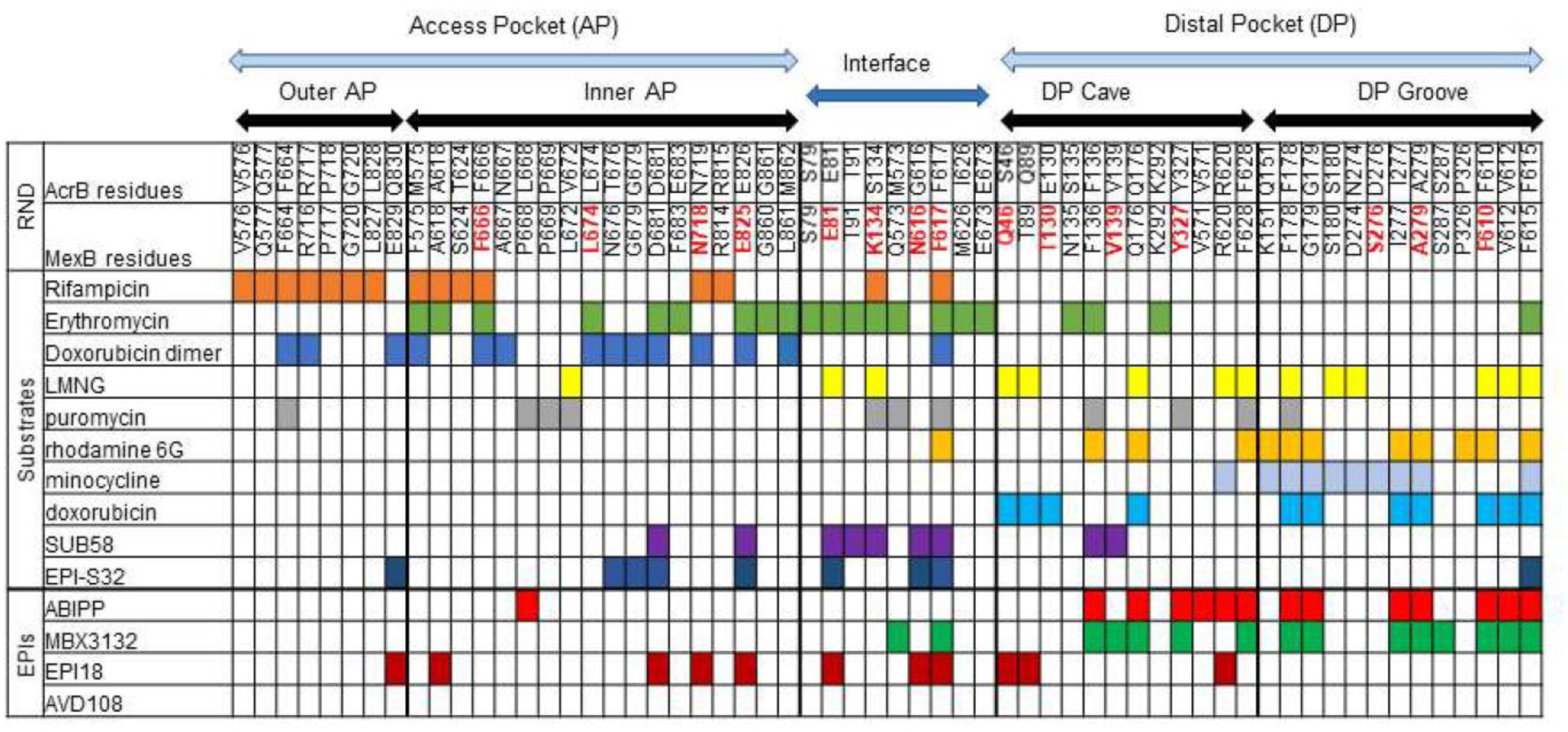
AcrB and MexB amino acid residues directly interacting with ligands from previous structural studies (modified from Kobylka et al.(16)) and the present investigation. Mutated amino acid residues of MexB are highlighted in red, all other residues are shown in black.

We first measured MICs of six antibiotics cefotaxime, levofloxacin, ciprofloxacin, novobiocin, chloramphenicol and trimethoprim, the known substrates of MexB (**Table 2**). All mutated MexB variants were expressed in *P. aeruginosa* PA2859 (ΔmexB ΔmexCD ΔmexYX) and its hyperporinated PA2859(Pore) derivative lacking the outer membrane barrier at comparable levels (**Figure S6**). However, the protein expression of the F610C (DP Groove) and F617C (IF) variants was ∼70% and 40% lower than the expression of MexB WT, respectively, suggesting larger scale structural changes in these mutants (**Figure S6**).

**Table 2.** MICs of antibiotics in PA2859 and PA2859(Pore) strains carrying an empty vector, plasmid-borne MexB and mutational variants. Shaded cells are MICs ≥4 folds change compared to MexB WT.

| Plasmid | MexB region | CEF<sup>a</sup> |  | LFX |  | CIP |  | NOV |  | CMP |  | TMP |  |
| --- | --- | --- | --- | --- | --- | --- | --- | --- | --- | --- | --- | --- | --- |
|  |  | Pore<sup>b</sup> | No Pore | Pore | No Pore | Pore | No Pore | Pore | No Pore | Pore | No Pore | Pore | No Pore |
| pUCP22 | NA | 0.078 | 1.25 | 0.008 | 0.0625 | 0.004 | 0.008 | 2 | 32 | 0.25 | 1 | 1.56 | 3.125 |
| MexB WT | NA | 0.625 | 20 | 0.125 | 1 | 0.016 | 0.125 | 128 | 1024 | 2 | 32 | 100 | 400 |
| Q46A | Cave | 0.625 | >10 | 0.125 | 1 | 0.016 | 0.125 | 128 | 1024 | 2 | 16 | 100 | 200 |
| T130C | Cave | 0.625 | 5-10 | 0.063 | 0.25-0.5 | 0.016 | 0.03 | 64 | 256-512 | 1 | 8 | 50 | >200 |
| V139C | Cave | 0.313 | 2.5 | 0.063 | 0.25 | 0.016 | 0.03 | 64 | 128 | 2 | 4 | 50 | 100 |
| V177A | Cave | 0.313 | 10 | 0.063 | 0.5-1 | 0.016 | 0.0625 | 64 | 1024 | 2 | 32 | 50 | >200 |
| Y327C | Cave | 0.313 | 10 | 0.125 | 0.5 | 0.016 | 0.063 | 64 | 128 | 2 | 16 | 100 | 200 |
| S276C | Groove | 0.625 | >10 | 0.125 | 1 | 0.031 | 0.063-0.125 | 128 | 1024 | 2 | 16-32 | 100 | >200 |
| A279V | Groove | 0.313 | 10-20 | 0.125 | 0.5 | 0.031 | 0.0625 | 128 | 128-256 | 2 | 32 | 100 | 200 |
| F610C | Groove | 0.625 | 2.5 | 0.031 | 0.125 | 0.004 | 0.016 | 16 | 128 | 1 | 2 | 3.125 | 25 |
| E81C | IF | 0.313 | >10 | 0.063 | 0.5-1 | 0.008 | 0.063 | 64 | 512 | 1 | 16-32 | 50 | >200 |
| K134C | IF | 0.156 | 5 | 0.125 | 1 | 0.016 | 0.125 | 64 | 512 | 2 | 32 | 50 | 200 |
| F617C | IF | 0.078 | 2.5 | 0.016 | 0.25 | 0.004 | 0.031 | 16 | 256 | 0.5 | 2 | 6.25 | 50 |
| N616A | IF | 0.625 | >10 | 0.063 | 1 | 0.016 | 0.063-0.125 | 128 | 512-1024 | 2 | 16 | 50 | >200 |
| F666C | Inner | 0.078 | 1.25 | 0.016 | 0.063 | 0.004 | 0.016 | 16 | 128 | 0.25 | 1 | 6.25 | 25 |
| V671A | Inner | 0.313 | 20 | 0.063 | 1 | 0.016 | 0.125 | 64 | 1024 | 1 | 16 | 50 | 200 |
| L674A | Inner | 0.625 | >10 | 0.125 | 1 | 0.016 | 0.125 | 128 | 1024 | 2 | 16 | 50 | >200 |
| N718C | Inner | 0.313 | 10 | 0.063 | 0.5 | 0.016 | 0.063 | 64 | 1024 | 1 | 8 | 50 | 100 |
| E825A | Inner | 0.625 | >10 | 0.125 | 1 | 0.016 | 0.125 | 128 | 1024 | 2 | 16 | 100 | >200 |

<sup>a</sup>, CEF -ceftaxime, LFX- levofloxacin, CIP- ciprofloxacin, NOV- novobiocin, CMP -chloramphenicol, TMP- trimethoprim.
<sup>b</sup>, Pore – PA2859(Pore), No Pore – PA2859.

MexB was the most efficient in the protection against novobiocin (NOV) and trimethoprim (TMP) (**Table 2**), with 32- to 64-fold increase in MICs independent of the outer membrane hyperporination. The pump was the least efficient against ciprofloxacin (CIP) producing a 16- and 4-fold change in MICs in PA2859 and PA2859(Pore), respectively. All constructed mutants were functional at least partially as judged from the complementation of the drug susceptible phenotype of PA2859 and its hyperporinated PA2859(Pore) derivative (**Table 2**).

Among the mutants, substitutions in the Inner AP residues F666 and N718 negatively affected the activity of MexB albeit to a different extent. Cells producing MexB F666C were hypersusceptible to all antibiotics, suggesting a non-specific effect. In contrast, MexB N718C was less effective only against chloramphenicol (CHL) (**Table 2**). Neither F666 nor N718 were among the top predictors but N718 interacted indirectly with EPI18 in the MD simulations (**Figure 5**).

The substitutions in the IF residues K134C and F617C demonstrated similar trend, with substitution in the aromatic F617C functionally defective likely due to non-specific changes and the loss of a positive charge in K134C selectively impaired against cefotaxime (CEF). K134 is a top predictor for MexB substrates and in the MD simulations, this residue formed water-mediated contacts with SUB58 and EPI-S32, but not EPI18 (**Figure 5**). F617C interacts directly with all three compounds, but these interactions are not predictive for substrates and inhibitors and the low expression level (**Figure S6**) suggests overall stability defects of this MexB variant.

The DP Cave substitutions T130C, V139C and Y327C also had specific and non-specific effects. The T130C substitution reduced the MICs of CEF, LFX, CIP, NOV and CHL by 4-fold, the Y327C substitution reduced the MIC of only NOV, whereas V139C showed a reduced activity against all tested antibiotics (**Table 2**). The V139C and Y327C residues are the top two predictors for MexB EPIs. These residues did not interact with the SUB58, EPI-S32 and EPI18 in MD simulations. However, the compounds interacted with the surrounding residues. For instance, T130 is located near V133, which interacts indirectly with EPI-S32 (13% of H-bridges persistence) (**Figure 5**).

Finally, the substitutions in DP Groove A279V and F610C followed the same trend in that A279V specifically reduced the MICs of NOV, whereas the replacement of the aromatic ring in F610C variant led to non-specific defects against all antibiotics. Neither one of these two residues contacted directly or indirectly the compounds in MD simulations. In addition, the lower expression of F610C variant could contribute to the compromised function.

The effect of all other substitutions was within the two-fold change in MICs, which is not significant for the two-fold dilution method used in this study. Thus, interactions of substrates within the Inner AP, the IF and the DP Cave and Groove of MexB are important for their recognition and transport. The specificity of interactions is apparently defined by non-hydrophobic amino acid side chains. The five top predictors of MexB substrates (K134 and V612) and inhibitors (V139, F178 and Y327) are functionally important.

### Both substrates and inhibitors potentiate activities of other antibiotics, albeit in ligand-and antibiotic-specific manner

The EPI activities can be assessed using two assays: (1) the bacterial growth-dependent assay that measures the ability of compounds to reduce the MIC of an antibiotic by at least four-fold (minimal potentiating concentration, MPC_4_) and (2) the inhibition of efflux of a fluorescent probe Hoechst 33342 (Hoechst) in non-growing bacterial cells. SUB58, EPI-S32 and EPI18 have only weak antibacterial activities (**Table S2**). We next determined MPC_4_ values for these three compounds in the presence of the best MexB substrates NOV and TMP. The experiments were carried out in PA2859(Pore) cells to reduce the contribution of the outer membrane barrier. Comparison of the MPC_4_ values determined for the three compounds showed that their activities are both antibiotic-and MexB variant-specific (**Table 3**). However, mutations in MexB could potentially affect either the translocation route of an EPI and/or an antibiotic or the inhibition route of an EPI. A comparison with changes in MICs could lead to distinguishing between these two possibilities.

**Table 3.** MPC_4_ of SUB58, EPI18, and EPI-S32 for novobiocin and trimethoprim.

| Plasmids<sup>a</sup> | MexB region | SUB58 | EPI-S32 |  | EPI18 |  |
| --- | --- | --- | --- | --- | --- | --- |
|  |  | NOV | NOV | TMP | NOV | TMP |
| **Vector** | NA | 25 | 6.25 | 25 | 3.1 | 6.3 |
| **MexB** | NA | 12.5 | 0.8 | 0.8 | 0.8 | 0.4 |
| **Q46A** | Cave | 12.5-25 | 0.4-0.8 | 0.2 | 0.8 | 0.8 |
| **T130C** | Cave | 12.5 | 0.4 | 0.2-0.4 | 0.8 | 0.8 |
| **V139C** | Cave | 12.5 | 0.2-0.4 | 0.2 | 0.8 | 0.8 |
| **V177A** | Cave | 12.5 | 0.8 | 0.4 | 0.8 | 0.8 |
| **Y327C** | Cave | 12.5-25 | 0.2-0.4 | 0.4 | 0.8 | 0.8-1.6 |
| **S276C** | Groove | 12.5-25 | 0.8-1.6 | 0.4 | 0.8-1.6 | 0.8-1.6 |
| **A279V** | Groove | 12.5-25 | 0.4 | 0.2 | 1.6 | 0.8-1.6 |
| **F610C** | Groove | 3.1 | 0.2 | 0.8 | 0.8-1.6 | 1.6 |
| **E81C** | IF | 25 | 3.13 | 0.4-0.8 | 1.6 | 0.8 |
| **K134C** | IF | 6.25-12.5 | 0.1-0.2 | 0.4-0.8 | 0.8 | 0.4 |
| **F617C** | IF | 3.1 | 0.2 | 0.8 | 0.8 | 0.8-1.6 |
| **N616A** | IF | 12.5 | 0.4-0.8 | 0.4 | 0.4-0.8 | 0.8 |
| **F666C** | Inner | 6.25-12.5 | 0.4 | 0.8-1.6 | 0.8-1.6 | 1.6 |
| **V671A** | Inner | 25 | 3.13 | 0.4-0.8 | 1.6 | 0.8 |
| **L674A** | Inner | 12.5 | 0.4-0.8 | 0.4-0.8 | 0.8 | 0.8 |
| **N718C** | Inner | 6.25 | 0.2-0.4 | 0.2 | 0.8 | 0.4 |
| **E825A** | Inner | 25 | 0.8 | 0.2-0.4 | 0.8 | 0.8 |

<sup>a</sup>, all measurements were done in PA2859(Pore) cells. Values shaded in green are with the decreased MPC<sub>4</sub> values and those shaded in red are with the increased MPC<sub>4</sub> values.

SUB58 potentiated the antibacterial activity of NOV but not the activity of TMP. The comparison of MPC_4_ of SUB58 and MICs of NOV showed that the MexB F610C and F617C variants that are the least effective against NOV are also the most susceptible to the potentiating activity of SUB58 (**Table 3**). Thus, the substrates NOV and SUB58, but not TMP, apparently compete for the same sites in MexB and the observed potentiation of NOV is due to competitive inhibition.

In contrast, EPI18 potentiated the activities of both NOV and TMP (**Table 3**). The potentiation of NOV was not sensitive to substitutions in MexB, suggesting that none of these substitutions is critical for the potentiating activity of EPI18. Interestingly, in the combination with TMP, higher concentrations of EPI18 were needed to reduce the MIC of TMP in cells producing MexB S276C, A279V and F610C (all in DP Groove), Y327C (DP Cave), F617C (IF) and F666C (Inner AP) variants, suggesting that these MexB variants were resistant to the inhibitory activity of EPI18, but only toward TMP not NOV.

Finally, like EPI18, EPI-S32 potentiated activities of both antibiotics but the effect of MexB substitutions was unique for this ligand. In the potentiation of the NOV activity, the MPC_4_ values of EPI-S32 were reduced in cells producing MexB with substitutions in all binding regions except the outer AP (**Table 3**), suggesting that all these variants are more sensitive to inhibitory activity of EPI-S32. Since V139C, F610C and F617C variants were less active against all substrates, their sensitivity to EPI-S32 could be non-specific. Surprisingly, E81C (IF) and V671A (Inner AP) became more resistant to this compound as seen from the MPC_4_ values and in the checkerboard assay (**Figure S7**). The profile of EPI-S32 and TMP combination was different. Substitutions Q46A, T130C and V139C, all in DP Cave, A279C in DP Groove, and N718C and E825C in Inner AP made MexB more sensitive to the potentiation of TMP by EPI-S32 (**Table 3**).

We also analyzed the activities of SUB58, EPI-S32, and EPI18 in the Hoechst efflux inhibition assay (**Figure 7**). In this assay, PA2859(Pore) cells producing MexB variants were pre-treated with increasing concentrations of the Rempex compounds and the intracellular accumulation of Hoechst was analyzed by following dye fluorescence in real-time. We found that in cells producing MexB WT, EPI18 efficiently inhibited efflux of Hoechst, as seen from increasing Hoechst fluorescence in cells treated with increasing concentrations of the compound (**Figure 7A**). In contrast, neither SUB58 nor EPI-S32 were able to inhibit efflux of Hoechst (**Figure 7B-C**). Hence, the mechanisms responsible for the potentiation of antibiotic activities and Hoechst efflux vary for the three compounds. In agreement, unlike with MPC_4_ measurement, cells producing the functionally compromised F610C, F617C and F666C variants as well as A279V, V671A and N718C were more sensitive to the EPI18 inhibition, as seen from the higher rates of Hoechst accumulation in these cells. In contrast, S276C (DP Groove) was resistant to EPI18 (**Figure 7D**).

**Figure 7.**
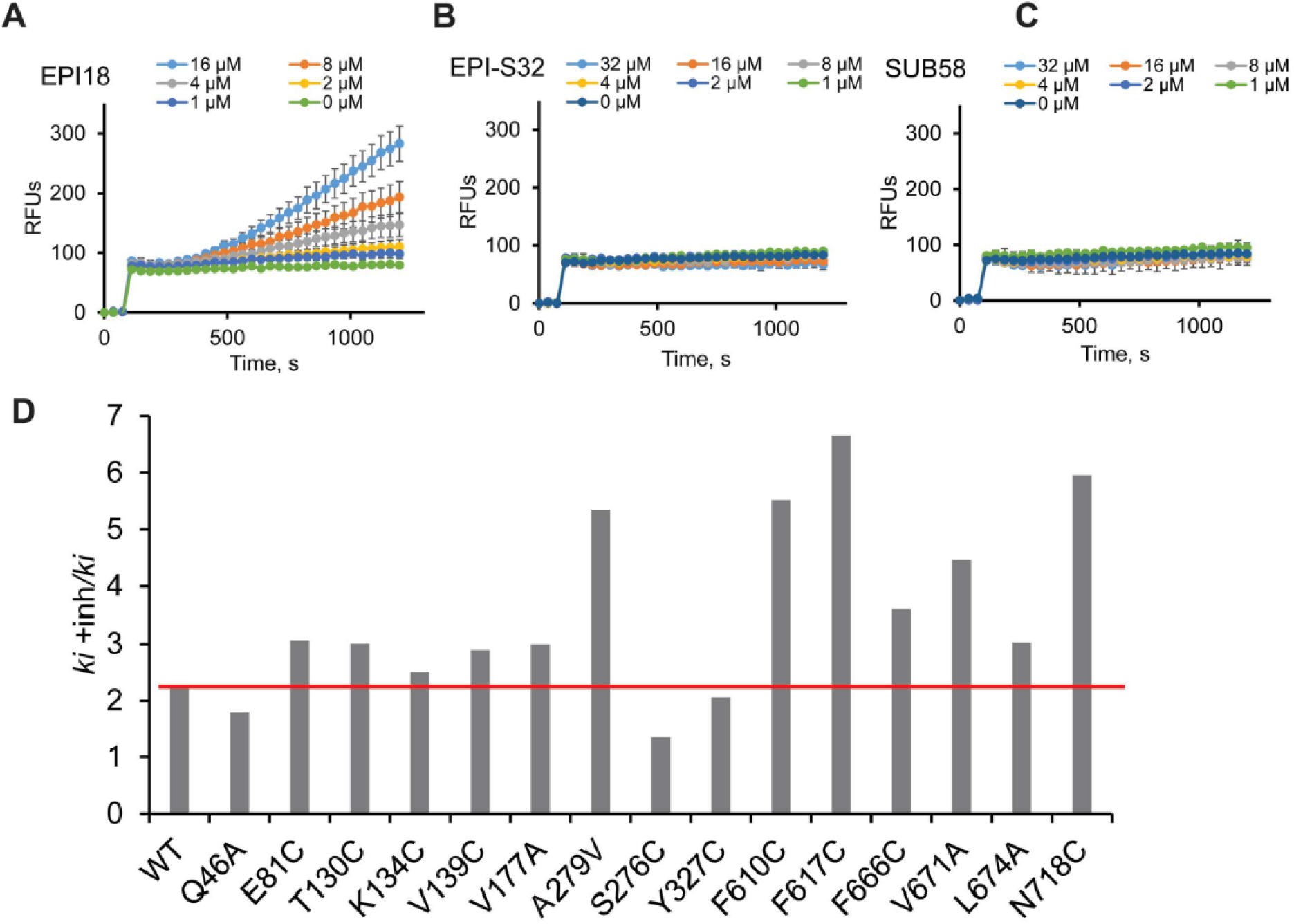
**Inhibition of MexB-dependent efflux of Hoechst**. PA2859(Pore) cells producing MexB WT were pre-incubated with increasing concentrations EPI18 (**A**), EPI-S32 (**B**) or SUB58 (**C**) for 15 min and then Hoechst was added to the final concentration 4 μM. The fluorescence was recorded in real-time. Error bars SD (n=3). **D.** The same as **A** but PA2859(Pore) cells producing the indicated MexB variants were pre-incubated without and with 8 μM of EPI18 and then Hoechst was added to the final concentration 4 μM. The kinetics of fluorescence change was analyzed in real-time. Kinetic curves were fitted to extract initial rates (*ki*) of Hoechst uptake. The ratios of initial rates in the presence (*ki*+inh) and absence (*ki*) of 8 μM EPI18 are shown. The red line shows the level of MexB WT.

Taken together, these results agree with the computational results and ML analyses that the substrates and inhibitors of MexB prefer different binding sites in MexB. Only in the case of EPI18, substitutions in the DP Groove and DP Cave of MexB lead to resistance against inhibition, whereas substitutions in the IF and Inner AP enable resistance against both EPI18 and EPI-S32. Several MexB substitutions in the Inner AP, IF and DP Cave that reduce MICs of antibiotic substrates also reduce MPC_4_ of Rempex compounds, whereas the substitutions in the DP Groove are associated only with efflux inhibitory activities of EPI-S32 and EPI18.

## Discussion

The increasing spread of antibiotic resistance in clinics demands new antimicrobial compounds. One of the most attractive and desirable features of antimicrobial compounds is the ability to avoid efflux. At the same time, identifying molecular determinants enabling the conversion of substrates to efflux inhibitors or avoiders is of paramount importance. Starting from the properties of Rempex compounds, the previously developed ML model was able to predict these properties in compounds that are structurally different from the originating ones and the calculated affinities to the MexB main binding sites as well as contacts with specific residues in these sites were among the top predictors (28). To gain further mechanistic and functional significance of the top predictive MexB residues discriminating between efflux substrates, avoiders, and inhibitors, in this work we combined ML analyses of docking results, MD simulations, and site-directed mutagenesis experiments. We focused on a series of 206 Rempex compounds representing the entire spectrum of ligand interactions with MexB starting from poor substrates/avoiders and all the way to excellent substrates or inhibitors.

The ensemble docking calculations indicate that this class of compounds bind preferentially in proximity to the channel 2 entrance (CH2), which is accessible from the periplasm and believed to be used by large antibiotics such as erythromycin and rifampicin (**Figure 6**)(48). In AcrB crystal structure, erythromycin was bound in the Inner AP, whereas rifampicin contacts residues in both the Inner and the Outer AP. The PC analysis of MexB contacts with the Rempex compounds derived from docking, showed that the Outer AP region occupies a very tight space on the PCA plot and in this respect, it is different from other ligand-binding regions, the residues of which are broadly spread on the plot and form intermixed clusters (**Figure 1**). Further inspection of the persistence of direct interactions of compounds calculated from MD simulations (**Table S3**) revealed a high number of contacts with the patch of negatively charged residues of the Outer AP of MexB that attracts the positively charged amino groups of Rempex compounds. This initial affinity to the Outer AP might be important for the high efficiency of efflux for good substrates and high efficiency of inhibition for EPIs, because AVD108 was found to have the lowest number of contacts with this region (**Figure 5**). In addition, the detailed analysis of water-mediated interactions suggests a prominent role of the solvent in all compounds except AVD108.

Some negatively charged residues of the Outer AP appear to interact with all representative compounds, whereas others are compound-specific. For instance, D566 in the Outer AP interacts with all four compounds, while the interaction with D568 seems to be unique for SUB58. Among the AP site residues that were mutated in this study, F666 appears to directly contact large antibiotics (**Figure 6**) and the substitution to cysteine likely changes the geometry of these interactions as seen from the drop in the MICs of all antibiotics in cells producing MexB F666C and an increase in MPC_4_ of EPI18 in combination with TMP (**Tables 2-3**). The N718C substitution on the other hand, is closer to the IF and leads to compound-specific changes in MIC and MPC_4_ values as well as the Hoechst efflux assay.

The persistence and the number of contacts with other binding regions in MexB appear to be compound-specific, so that even small chemical modifications can give rise to different interaction patterns. This compound-specific behaviour is seen from the dispersion of the IF and DP residues on the PCA plot (**Figure 1B**). Nevertheless, certain interactions are specifically associated with the substrate translocation and the inhibition routes, respectively. Among the top ML predictors of MexB substrates, Q46, K134 and R620 are located at or near the IF that separates the AP from the DP and likely reflect the importance of the IF interactions in the translocation of substrates, in addition to those in the DP Groove and Cave. Importantly, all top side chain predictors of MexB substrates are non-aromatic and predominantly (5 out of 6) are either charged or polar. On the contrary, the residues in the DP Cave and Groove of MexB predictive of EPIs are all aromatic and hydrophobic, in agreement with structural findings of the importance of aromatic residues for the inhibition of MexB and related pumps (18)(41). Furthermore, the mutagenesis data provided a strong support for the functional distinction between the aromatic and polar residues in the DP of MexB (**Tables 2**). The substitutions in the aromatic residues, as perhaps binding of EPIs to these residues, lead to broad non-specific changes in the activity or even inactivation of MexB, whereas substitutions in polar and charged residues affect specific ligands.

MD simulations showed that all Rempex compounds expose their molecular portions containing aromatic rings towards the interior of the AP and are attracted toward the Inner AP and the IF by hydrophobic forces. All three ligands SUB58, EPI-S32 and EPI18 interact with multiple residues in the IF region that are also engaged by other ligands (**Figure 6**). To get access into the DP, all compounds must squeeze through the switch loop (residues 613-623), which seems to impose the restriction on the size of ligands (49). MD simulations suggested that N616 is important for the interaction with EPI18 but not with SUB58 (**Figure 4**). The N616A substitution, however, had no effect on antibacterial activities of antibiotics and inhibitory properties of EPI18. The substitution F617C in this loop, on the other hand, reduced efflux efficiency against all tested antibiotics as judged from the low MIC values of antibiotics (**Table 2**). Hence, in agreement with previous studies, the conformation of the loop and its interactions with substrates are important for ligand translocation (50)(51). Surprisingly, the same F617C substitution makes MexB more resistant to the action of EPI18 in combination with TMP, but not with NOV (**Table 3**). Mutations in MexB could potentially affect either the translocation route of an EPI and/or an antibiotic or the inhibition route of an EPI. Since TMP is one of the smallest and most hydrophilic ligands of MexB (**Figure 2**), the substitutions in the switch loop are unlikely to hinder the TMP translocation and the observed resistance is likely due to problems with the translocation of EPI18 by MexB F617C variant. In contrast, the potentiating effect in the inhibition of efflux of Hoechst by EPI18 or the lower MPC_4_ of EPI-S32 and SUB58 in combination with NOV, are likely due to the structural role of F617 in the translocation of all these ligands. Accordingly, MD simulations showed a network of both direct and indirect interactions between EPI-S32 and EPI18 and F617 (14% and 18% of persistence of direct interactions for EPI-S32 and EPI18, respectively, and 14% of indirect interactions in EPI18).

Similarly in the DP, all ligands interact directly or indirectly with multiple residues, some of which are the top side chain predictors of substrates and EPIs. In agreement, several mutations in the Cave and the Groove region impacted both the translocation and the inhibition routes (**Tables 2-3**, **Figure 7**). For example, the Y327C mutation in MexB makes cells more susceptible to antibacterial activity of NOV and to the inhibitory activity of EPI-S32 in combination with NOV. On the other hand, the same substitution makes cells more resistant to the EPI18 inhibition in the combination with TMP, likely by affecting the inhibition route. The importance of this residue was also demonstrated for some MexB homologs, and as we found for MexB, the effect of substitutions was ligand specific (reviewed in Kobylka et al. (16)). For example, *E. coli* AcrB with the Y327A substitution was reported to be more susceptible to minocycline, rhodamine-6G, tetraphenylphosphonium, LVX and CHL (17), while in *A. baumannii* AdeB with the same mutation increased the resistance towards LVX and CHL, suggesting more efficient efflux (17).

Taken together, our findings suggest that steric and conformational changes associated with mutations in the Groove and Cave of the DP of MexB and other RND pumps, affect differently the interaction network driving substrate translocation and inhibition of the pump. The ML and molecular-level description of the interaction between the Rempex compounds and MexB show that the translocation and the inhibition routes are common in the AP and split in the IF and DP with some compounds engaging the right combination of residues in the IF and the DP Cave to be effectively expelled into the exit funnel, whereas others engage aromatic residues in the DP Groove and DP Cave that slow down or hinder the next step in the transporter cycle. These findings suggest that all MexB ligands fit into this substrate-inhibitor spectrum depending on their physico-chemical structures and properties. The ML models and MD simulations identify different aspects of ligand-transporter interactions, with the former picking rare interactions important for distinguishing between inhibitor vs. non-inhibitor or substrate vs. avoider. In contrast, MD simulations capture residues important for recognition of all ligands and their subsequent translocation. Our results do not exclude the possibility that inhibition of efflux is achieved when the AP (or IF) and DP are both occupied with high-affinity ligands in the same or the two adjacent Loose and Tight protomers of MexB. The identified residues and pathways can be further used for the rational design of new EPIs and effective efflux avoiders by enhancing interactions that were found associated with EPIs and discouraging those important for substrate translocation, respectively.

## METHODS

### Ensemble docking to MexB

Molecular docking calculations were performed with the software AutoDock VINA (52), implementing a stochastic global optimization approach. The program was used with default settings but for the exhaustiveness (giving a measure of the exhaustiveness of the local search) which was set to 1024 (default 8). Protein and ligand input files were prepared with AutoDock Tools (53). Flexibility of both docking partners was considered indirectly by using ensembles of conformations. For each compound we used 10 different cluster representatives extracted from MD simulations in explicit water solution, while for MexB we considered 6 conformations including available X-ray structures (PDB Ids: 2V50 (19), 3W9J, 3W9I (18)), and MD snapshots (15). For each docking run we retained the top ten docking poses. We performed two sets of guided docking runs into AP_L_ and DP_T_. In each case the search of poses was performed within a cubic volume of 40x40x40 Å^3^.

### Machine learning

Machine Learning analysis was performed using the Statistics and Machine Learning toolbox of Matlab. In lieu of descriptors, we used the per-residue interaction counts generated for each compound in the previous study (28). To reduce the overfitting during classification, the interaction counts were binned on a common scale into a total of 16 bins 20 counts each. The experimental efflux or inhibition ratios were transformed to a logarithmic scale and imputed by approximating all ratios smaller than 1 with 1. The experimental data were then balanced by the introduction of statistical weights that were inversely related to the bin counts of the compounds after the logarithms of the ratios were binned into 10 equal intervals. A “random forest” ensemble of 100,000 regression trees was then assembled using bootstrap aggregation and the curvature test for descriptor selection (54). The descriptors were then ranked in the descending order according to their associated misclassification cost.

### Molecular dynamics

The initial coordinates of the MexB/Rempex complexes were embedded in a pre-equilibrated 1-palmitoyl-2-oleoyl-*sn*-glycero-3-phosphoethanolamine (POPE) bilayer patch. The embedding of the complex into the POPE bilayer was performed as detailed previously (15). Each system was then immersed in a box containing TIP3P water molecules and an adequate number of K^+^ counterions to neutralize the negative net charge of the system. An osmolarity value of 0.15 M was reached by adding an appropriate number of K^+^/Cl^−^. The *ff*14SB versions of the Amber force field and lipid14 were adopted for MexB and the POPE bilayer, respectively. The GAFF2 (55) parameters adopted for the ligands were derived as reported previously (56). The systems were minimized with a combination of the steepest descent and conjugate gradient methods gradually releasing positional restraints applied. The systems were heated from 0 to 310 K in two steps: a 1 ns heating from 0 to 100 K in a canonical ensemble (NVT) followed by a 5 ns heating to reach 310 K in an isothermal–isobaric ensemble (NPT) while applying positional restraints along the *Z* axis on the phosphorous heads of lipids molecules, allowing the merging of the membrane. Multiple equilibration steps of 500 ps each until the stabilization of the box dimensions were performed in the NPT ensemble. A Langevin thermostat using a collision frequency of 1 ps^∓1^ and a Berendsen isotropic barostat maintained a constant temperature and an average pressure of 1 Atm, respectively. A time step of 2 fs was used during the equilibration protocol. The MD simulations were carried out using the PMEMD module of Amber14 with a time-step of 4 fs in the NVT ensemble after applying the hydrogen mass repartitioning. Coordinates were saved every 100 ps. Long-range electrostatic interactions were calculated using the particle mesh Ewald method with a cut-off of 9 Å.

### MD analyses

The CPPTRAJ(57) module of AmberTools14 was used to (1) cluster the trajectories according to the RMSDs of ligands, using a hierarchical algorithm (58), (2) compute the MexB-ligands contacts during the simulations, and (3) identify the water-mediated H-bridges. In details, for the points (2) and (3), both the ligands and the side chains residues of MexB were divided into pharmacophoric groups, and the distances/interactions were computed between compatible pharmacophoric types (e.g., distance between H-bond donor and acceptor, positively and negatively charged groups, π-π between aromatic rings, etc) (see **Figure S9**).

### Site-directed mutagenesis

The gene encoding MexB was PCR amplified from *P. aeruginosa* PAO1 strain and cloned into pUCP22 vector using standard protocols. Site-directed mutagenesis was carried out using the Quick-Change protocol. All substitutions were verified by DNA sequencing (Oklahoma Medical Research Foundation DNA sequencing facility). For expression and functional analyses, plasmids producing MexB variants were transformed into the parental strain PA2859 (ΔmexB ΔmexCD ΔmexXY) strain and its hyperporinated derivative PA2859(Pore). The hyperporinated PA2859(Pore) strain was constructed as described previously (59).

### Antibiotic sensitivity/susceptibility assay

MIC of selected antibiotics against MexB mutants was measured using a two-fold serial dilution broth assay as described previously (28). Briefly, overnight grown cells were sub-cultured 1:100 into fresh LB media until OD_600_∼0.2 followed by induction with 0.1 mM IPTG for 3 hours. MIC plate was set up with 2-fold dilution of compound concentration in 100 μl of LB broth per well. A positive and negative control well with cells only and media only were included in each plate, respectively. Cells 10^3^·10^4^ were added to each well except negative control wells. Plates were incubated at 37 °C for 18-24 hours and OD_600_ was measured using a Spark 10M microplate reader (Tecan) and the data was analyzed and expressed as MIC.

### Hoechst efflux inhibition assay

Substrate (Hoechst) efflux assay was performed in a temperature-controlled microplate reader (Tecan Spark 10M) in fluorescence mode as described before (28). Briefly, overnight grown cells were sub-cultured into a fresh LB medium and grown at 37 °C to an optical density at 600 nm (OD_600_) of ∼0.2 followed by induction with 0.1 mM IPTG for another ∼3 hours. The cells in the exponential phase were collected by centrifugation at 4,000 rpm for 10 min at room temperature (RT). The cells were washed using HMG buffer (50 mM HEPES-KOH buffer pH 7.0, 1 mM magnesium sulfate and 0.4 mM glucose) and resuspended in HMG buffer to an OD_600_ of ∼1.0, at RT. Increasing concentrations of ligands were added to the cells and after 10 min incubation at RT, Hoechst was added to the final concentration 4 μM. Fluorescence of Hoechst was monitored in real time at λ_ex_ = 350 nm and λ_em_ = 450 nm.

### Growth-dependent inhibition/potentiation assays

The potentiation activity was measured for an EPI18, EPI-S32 and SUB58 as described previously (28). MPC_4_ is defined as the concentration of the EPI/compound that reduces the MIC of antibiotic by four-fold. The MPC_4_ values for each compound were determined for each mutant/control based on non-visible growth of cells at that specific concentration of the compound.

## Acknowledgements

This work was supported by NIAID/NIH grant number R01AI136799. S.G., G.M., and P.R. gratefully acknowledge the Health Extended ALliance for Innovative Therapies, Advanced Lab-research, and Integrated Approaches of Precision Medicine partnership (HEAL ITALIA), foundation by the Italian Ministry of University and Research, PNRR, mission 4, component 2, investment 1.3, project number PE00000019 (University of Cagliari). A.V.V gratefully acknowledges the “One Health Basic and Translational Research Actions addressing Unmet Needs on Emerging Infectious Diseases (INF-ACT)” foundation by the Italian Ministry of University and Research, PNRR, mission 4, component 2, investment 1.3, project number PE00000007 (University of Cagliari).

## Authors’ contributions

S.G., G.M., A.B., E.M., and A.V.V. carried out MD simulations and docking experiments with MexB. J.M. and I.V.L. constructed MexB mutant variants and carried out analyses of biological activities of Rempex compounds. C.B. performed statistical analyses and generated machine learning models. P.R., V.V.R. and H.I.Z. designed the studies, analyzed the data, and wrote the manuscript. All authors contributed to writing of the manuscript.

## REFERENCES

1. Fujiwara M, Yamasaki S, Morita Y, Nishino K. 2022. Evaluation of efflux pump inhibitors of MexAB- or MexXY-OprM in Pseudomonas aeruginosa using nucleic acid dyes. J Infect Chemother 28:595–601.

2. Murray CJ, Ikuta KS, Sharara F, Swetschinski L, Robles Aguilar G, Gray A, Han C, Bisignano C, Rao P, Wool E, Johnson SC, Browne AJ, Chipeta MG, Fell F, Hackett S, Haines-Woodhouse G, Kashef Hamadani BH, Kumaran EAP, McManigal B, Agarwal R, Akech S, Albertson S, Amuasi J, Andrews J, Aravkin A, Ashley E, Bailey F, Baker S, Basnyat B, Bekker A, Bender R, Bethou A, Bielicki J, Boonkasidecha S, Bukosia J, Carvalheiro C, Castañeda-Orjuela C, Chansamouth V, Chaurasia S, Chiurchiù S, Chowdhury F, Cook AJ, Cooper B, Cressey TR, Criollo-Mora E, Cunningham M, Darboe S, Day NPJ, De Luca M, Dokova K, Dramowski A, Dunachie SJ, Eckmanns T, Eibach D, Emami A, Feasey N, Fisher-Pearson N, Forrest K, Garrett D, Gastmeier P, Giref AZ, Greer RC, Gupta V, Haller S, Haselbeck A, Hay SI, Holm M, Hopkins S, Iregbu KC, Jacobs J, Jarovsky D, Javanmardi F, Khorana M, Kissoon N, Kobeissi E, Kostyanev T, Krapp F, Krumkamp R, Kumar A, Kyu HH, Lim C, Limmathurotsakul D, Loftus MJ, Lunn M, Ma J, Mturi N, Munera-Huertas T, Musicha P, Mussi-Pinhata MM, Nakamura T, Nanavati R, Nangia S, Newton P, Ngoun C, Novotney A, Nwakanma D, Obiero CW, Olivas-Martinez A, Olliaro P, Ooko E, Ortiz-Brizuela E, Peleg AY, Perrone C, Plakkal N, Ponce-de-Leon A, Raad M, Ramdin T, Riddell A, Roberts T, Robotham JV, Roca A, Rudd KE, Russell N, Schnall J, Scott JAG, Shivamallappa M, Sifuentes-Osornio J, Steenkeste N, Stewardson AJ, Stoeva T, Tasak N, Thaiprakong A, Thwaites G, Turner C, Turner P, van Doorn HR, Velaphi S, Vongpradith A, Vu H, Walsh T, Waner S, Wangrangsimakul T, Wozniak T, Zheng P, Sartorius B, Lopez AD, Stergachis A, Moore C, Dolecek C, Naghavi M. 2022. Global burden of bacterial antimicrobial resistance in 2019: a systematic analysis. Lancet 399:629–655.

3. Rojas A, Palacios-Baena ZR, L Opez-Cort Es LE, Rodríguez-Ba J. 2019. Rates, predictors and mortality of community-onset bloodstream infections due to Pseudomonas aeruginosa: systematic review and meta-analysis. Clin Microbiol Infect 25:964.

4. Ma M, Lustig M, Salem M, Mengin-lecreulx D, Phan G, Broutin I. 2021. MexAB-OprM Efflux Pump Interaction with the Peptidoglycan of Escherichia coli and Pseudomonas aeruginosa. Int J Mol Sci 2021, Vol 22, Page 5328 22:5328.

5. Alav I, Kobylka J, Kuth MS, Pos KM, Picard M, Blair JMA, Bavro VN. 2021. Structure, Assembly, and Function of Tripartite Efflux and Type 1 Secretion Systems in Gram-Negative Bacteria. Chem Rev 121:5479–5596.

6. Klenotic PA, Moseng MA, Morgan CE, Yu EW. 2021. Structural and Functional Diversity of Resistance-Nodulation-Cell Division Transporters. Chem Rev 121:5378–5416.

7. Pang Z, Raudonis R, Glick BR, Lin TJ, Cheng Z. 2019. Antibiotic resistance in Pseudomonas aeruginosa: mechanisms and alternative therapeutic strategies. Biotechnol Adv 37:177–192.

8. Tsutsumi K, Yonehara R, Ishizaka-Ikeda E, Miyazaki N, Maeda S, Iwasaki K, Nakagawa A, Yamashita E. 2019. Structures of the wild-type MexAB–OprM tripartite pump reveal its complex formation and drug efflux mechanism. Nat Commun 10:1–10.

9. Zahedi bialvaei A, Rahbar M, Hamidi-Farahani R, Asgari A, Esmailkhani A, Mardani dashti Y, Soleiman-Meigooni S. 2021. Expression of RND efflux pumps mediated antibiotic resistance in Pseudomonas aeruginosa clinical strains. Microb Pathog 153:104789.

10. Zgurskaya HI, Malloci G, Chandar B, Vargiu A V., Ruggerone P. 2021. Bacterial efflux transporters’ polyspecificity - a gift and a curse? Curr Opin Microbiol 61:115–123.

11. Sakurai K, Yamasaki S, Nakao K, Nishino K, Yamaguchi A, Nakashima R. 2019. Crystal structures of multidrug efflux pump MexB bound with high-molecular-mass compounds. Sci Reports 2019 91 9:1–9.

12. Huang L, Wu C, Gao H, Xu C, Dai M, Huang L, Hao H, Wang X, Cheng G. 2022. Bacterial Multidrug Efflux Pumps at the Frontline of Antimicrobial Resistance: An Overview. Antibiotics 11:520.

13. Schulz R, Vargiu A V., Collu F, Kleinekathöfer U, Ruggerone P. 2010. Functional Rotation of the Transporter AcrB: Insights into Drug Extrusion from Simulations. PLOS Comput Biol 6:e1000806.

14. Du D, Wang-Kan X, Neuberger A, van Veen HW, Pos KM, Piddock LJV, Luisi BF. 2018. Multidrug efflux pumps: structure, function and regulation. Nat Rev Microbiol 16:523–539.

15. Ramaswamy VK, Vargiu A V., Malloci G, Dreier J, Ruggerone P. 2018. Molecular determinants of the promiscuity of MexB and MexY multidrug transporters of Pseudomonas aeruginosa. Front Microbiol 9:1144.

16. Kobylka J, Kuth MS, Müller RT, Geertsma ER, Pos KM. 2020. AcrB: a mean, keen, drug efflux machine. Ann N Y Acad Sci 1459:38–68.

17. Ornik-Cha A, Wilhelm J, Kobylka J, Sjuts H, Vargiu A V., Malloci G, Reitz J, Seybert A, Frangakis AS, Pos KM. 2021. Structural and functional analysis of the promiscuous AcrB and AdeB efflux pumps suggests different drug binding mechanisms. Nat Commun 12:1–14.

18. Nakashima R, Sakurai K, Yamasaki S, Hayashi K, Nagata C, Hoshino K, Onodera Y, Nishino K, Yamaguchi A. 2013. Structural basis for the inhibition of bacterial multidrug exporters. Nature 500:102–106.

19. Sennhauser G, Bukowska MA, Briand C, Grütter MG. 2009. Crystal structure of the multidrug exporter MexB from Pseudomonas aeruginosa. J Mol Biol 389:134–145.

20. Glavier M, Puvanendran D, Salvador D, Decossas M, Phan G, Garnier C, Frezza E, Cece Q, Schoehn G, Picard M, Taveau JC, Daury L, Broutin I, Lambert O. 2020. Antibiotic export by MexB multidrug efflux transporter is allosterically controlled by a MexA-OprM chaperone-like complex. Nat Commun 11:4948.

21. Eicher T, Cha HJ, Seeger MA, Brandstätter L, El-Delik J, Bohnert JA, Kern W V., Verrey F, Grütter MG, Diederichs K, Pos KM. 2012. Transport of drugs by the multidrug transporter AcrB involves an access and a deep binding pocket that are separated by a switch-loop. Proc Natl Acad Sci U S A 109:5687–5692.

22. Mangiaterra G, Laudadio E, Cometti M, Mobbili G, Minnelli C, Massaccesi L, Citterio B, Biavasco F, Galeazzi R. 2017. Inhibitors of multidrug efflux pumps of Pseudomonas aeruginosa from natural sources: An in silico high-throughput virtual screening and in vitro validation. Med Chem Res 26:414–430.

23. Zhao S, Adamiak JW, Bonifay V, Mehla J, Zgurskaya HI, Tan DS. 2020. Defining new chemical space for drug penetration into Gram-negative bacteria. Nat Chem Biol 16:1293–1302.

24. Cooper CJ, Krishnamoorthy G, Wolloscheck D, Walker JK, Rybenkov V V., Parks JM, Zgurskaya HI. 2018. Molecular Properties That Define the Activities of Antibiotics in Escherichia coli and Pseudomonas aeruginosa. ACS Infect Dis 4:1223–1234.

25. Cacciotto P, Ramaswamy VK, Malloci G, Ruggerone P, Vargiu A V. 2018. Molecular Modeling of Multidrug Properties of Resistance Nodulation Division (RND) Transporters. Methods Mol Biol 1700:179–219.

26. Atzori A, Malloci G, Prajapati JD, Basciu A, Bosin A, Kleinekathöfer U, Dreier J, Vargiu A V., Ruggerone P. 2019. Molecular Interactions of Cephalosporins with the Deep Binding Pocket of the RND Transporter AcrB. J Phys Chem B 123:4625–4635.

27. Vergalli J, Atzori A, Pajovic J, Dumont E, Malloci G, Masi M, Vargiu AV, Winterhalter M, Réfrégiers M, Ruggerone P, Pagès JM. 2020. The challenge of intracellular antibiotic accumulation, a function of fluoroquinolone influx versus bacterial efflux. Commun Biol 3:1–12.

28. Mehla J, Malloci G, Mansbach R, López CA, Tsivkovski R, Haynes K, Leus I V., Grindstaff SB, Cascella RH, D’cunha N, Herndon L, Hengartner NW, Margiotta E, Atzori A, Vargiu A V., Manrique PD, Walker JK, Lomovskaya O, Ruggerone P, Gnanakaran S, Rybenkov V V., Zgurskaya HI. 2021. Predictive rules of efflux inhibition and avoidance in pseudomonas aeruginosa. MBio 12:1–19.

29. Zgurskaya HI, Walker JK, Parks JM, Rybenkov V V. 2021. Multidrug Efflux Pumps and the Two-Faced Janus of Substrates and Inhibitors. Acc Chem Res 54:930–939.

30. Lomovskaya O, Warren MS, Lee A, Galazzo J, Fronko R, Lee M, Blais J, Cho D, Chamberland S, Renau T, Leger R, Hecker S, Watkins W, Hoshino K, Ishida H, Lee VJ. 2001. Identification and characterization of inhibitors of multidrug resistance efflux pumps in Pseudomonas aeruginosa: Novel agents for combination therapy. Antimicrob Agents Chemother 45:105–116.

31. Sjuts H, Vargiu A V., Kwasny SM, Nguyen ST, Kim HS, Ding X, Ornik AR, Ruggerone P, Bowlin TL, Nikaido H, Pos KM, Opperman TJ. 2016. Molecular basis for inhibition of AcrB multidrug efflux pump by novel and powerful pyranopyridine derivatives. Proc Natl Acad Sci U S A 113:3509– 3514.

32. Wang Y, Mowla R, Ji S, Guo L, De Barros Lopes MA, Jin C, Song D, Ma S, Venter H. 2018. Design, synthesis and biological activity evaluation of novel 4-subtituted 2-naphthamide derivatives as AcrB inhibitors. Eur J Med Chem 143:699–709.

33. Garvey MI, Rahman MM, Gibbons S, Piddock LJV. 2011. Medicinal plant extracts with efflux inhibitory activity against Gram-negative bacteria. Int J Antimicrob Agents 37:145–151.

34. Piddock LJV, Garvey MI, Rahman MM, Gibbons S. 2010. Natural and synthetic compounds such as trimethoprim behave as inhibitors of efflux in Gram-negative bacteria. J Antimicrob Chemother 65:1215–1223.

35. Plé C, Tam HK, Vieira Da Cruz A, Compagne N, Jiménez-Castellanos JC, Müller RT, Pradel E, Foong WE, Malloci G, Ballée A, Kirchner MA, Moshfegh P, Herledan A, Herrmann A, Deprez B, Willand N, Vargiu AV, Pos KM, Flipo M, Hartkoorn RC. 2022. Pyridylpiperazine-based allosteric inhibitors of RND-type multidrug efflux pumps. Nat Commun 2022 131 13:1–11.

36. Aron Z, Opperman TJ. 2018. The hydrophobic trap—the Achilles heel of RND efflux pumps. Res Microbiol 169:393–400.

37. Vergalli J, Chauvet H, Oliva F, Pajović J, Malloci G, Vargiu AV, Réfrégiers M, Ruggerone P, Pagès JM. 2022. A framework for dissecting affinities of multidrug efflux transporter AcrB to fluoroquinolones. Commun Biol 5:1–11.

38. Ciusa ML, Marshall RL, Ricci V, Stone JW, Piddock LJV. 2022. Absence, loss-of-function, or inhibition of Escherichia coli AcrB does not increase expression of other efflux pump genes supporting the discovery of AcrB inhibitors as antibiotic adjuvants. J Antimicrob Chemother 77:633–640.

39. Renau TE, Léger R, Filonova L, Flamme EM, Wang M, Yen R, Madsen D, Griffith D, Chamberland S, Dudley MN, Lee VJ, Lomovskaya O, Watkins WJ, Ohta T, Nakayama K, Ishida Y. 2003. Conformationally-restricted analogues of efflux pump inhibitors that potentiate the activity of levofloxacin in Pseudomonas aeruginosa. Bioorganic Med Chem Lett 13:2755–2758.

40. Vargiu AV, Ramaswamy VK, Malvacio I, Malloci G, Kleinekathöfer U, Ruggerone P. 2018. Water-mediated interactions enable smooth substrate transport in a bacterial efflux pump. Biochim Biophys Acta - Gen Subj 1862:836–845.

41. Vargiu A V., Ruggerone P, Opperman TJ, Nguyen ST, Nikaido H. 2014. Molecular mechanism of MBX2319 inhibition of Escherichia coli AcrB multidrug efflux pump and comparison with other inhibitors. Antimicrob Agents Chemother 58:6224–6234.

42. D’Cunha N, Moniruzzaman M, Haynes K, Malloci G, Cooper CJ, Margiotta E, Vargiu A V., Uddin MR, Leus I V., Cao F, Parks JM, Rybenkov V V., Ruggerone P, Zgurskaya HI, Walker JK. 2021. Mechanistic Duality of Bacterial Efflux Substrates and Inhibitors: Example of Simple Substituted Cinnamoyl and Naphthyl Amides. ACS Infect Dis 7:2650–2665.

43. Jollife IT, Cadima J. 2016. Principal component analysis: A review and recent developments. Philos Trans R Soc A Math Phys Eng Sci 374:2065.

44. Yamaguchi A, Nakashima R, Sakurai K. 2015. Structural basis of RND-type multidrug exporters. Front Microbiol 6:327.

45. Haloi N, Vasan AK, Geddes EJ, Prasanna A, Wen PC, Metcalf WW, Hergenrother PJ, Tajkhorshid E. 2021. Rationalizing the generation of broad spectrum antibiotics with the addition of a positive charge. Chem Sci 12:15028–15044.

46. Mao W, Warren MS, Black DS, Satou T, Murata T, Nishino T, Gotoh N, Lomovskaya O. 2002. On the mechanism of substrate specificity by resistance nodulation division (RND)-type multidrug resistance pumps: the large periplasmic loops of MexD from Pseudomonas aeruginosa are involved in substrate recognition. Mol Microbiol 46:889–901.

47. Tikhonova EB, Wang Q, Zgurskaya HI. 2002. Chimeric analysis of the multicomponent multidrug efflux transporters from gram-negative bacteria. J Bacteriol 184:6499–6507.

48. Zwama M, Yamasaki S, Nakashima R, Sakurai K, Nishino K, Yamaguchi A. 2018. Multiple entry pathways within the efflux transporter AcrB contribute to multidrug recognition. Nat Commun 9:124.

49. Müller RT, Travers T, Cha H jea, Phillips JL, Gnanakaran S, Pos KM. 2017. Switch Loop Flexibility Affects Substrate Transport of the AcrB Efflux Pump. J Mol Biol 429:3863–3874.

50. Bohnert JA, Schuster S, Seeger MA, Fähnrich E, Pos KM, Kern W V. 2008. Site-directed mutagenesis reveals putative substrate binding residues in the Escherichia coli RND efflux pump AcrB. J Bacteriol 190:8225–8229.

51. Vargiu A V., Collu F, Schulz R, Pos KM, Zacharias M, Kleinekathöfer U, Ruggerone P. 2011. Effect of the F610A mutation on substrate extrusion in the AcrB transporter: Explanation and rationale by molecular dynamics simulations. J Am Chem Soc 133:10704–10707.

52. Trott O, Olson AJ. 2010. AutoDock Vina: Improving the speed and accuracy of docking with a new scoring function, efficient optimization, and multithreading. J Comput Chem 31:455–461.

53. Morris GM, Ruth H, Lindstrom W, Sanner MF, Belew RK, Goodsell DS, Olson AJ. 2009. AutoDock4 and AutoDockTools4: Automated docking with selective receptor flexibility. J Comput Chem 30:2785–2791.

54. Loh WY, Shin YS. 1997. Split selection methods for classification trees. Stat Sin 7:815–840.

55. Wang J, Wolf RM, Caldwell JW, Kollman PA, Case DA. 2004. Development and testing of a general amber force field. J Comput Chem 25:1157–1174.

56. Gervasoni S, Malloci G, Bosin A, Vargiu A V., Zgurskaya HI, Ruggerone P. 2022. AB-DB: Force-Field parameters, MD trajectories, QM-based data, and Descriptors of Antimicrobials. Sci Data 2022 91 9:1–12.

57. Roe DR, Cheatham TE. 2013. PTRAJ and CPPTRAJ: Software for processing and analysis of molecular dynamics trajectory data. J Chem Theory Comput 9:3084–3095.

58. Shao J, Tanner SW, Thompson N, Cheatham TE. 2007. Clustering molecular dynamics trajectories: 1. Characterizing the performance of different clustering algorithms. J Chem Theory Comput 3:2312–2334.

59. Krishnamoorthy G, Leus I V., Weeks JW, Wolloscheck D, Rybenkov V V., Zgurskaya HI. 2017. Synergy between Active Efflux and Outer Membrane Diffusion Defines Rules of Antibiotic Permeation into Gram-Negative Bacteria. MBio 8:e01172–17.

