## Supplementary Information for "Molecular determinants of avoidance and inhibition of *Pseudomonas aeruginosa* MexB efflux pump"

**Figure S1.** Spatial distribution of MexB regions. Left: Architecture of the substrate binding site of MexB with its five main areas highlighted as following: Outer AP-red, Inner AP – green, Interface – blue, DP Groove – gold; DP Cave – cyan. Right: The switch loop is colored in blue and the residues lining the hydrophobic trap are reported as red stick.

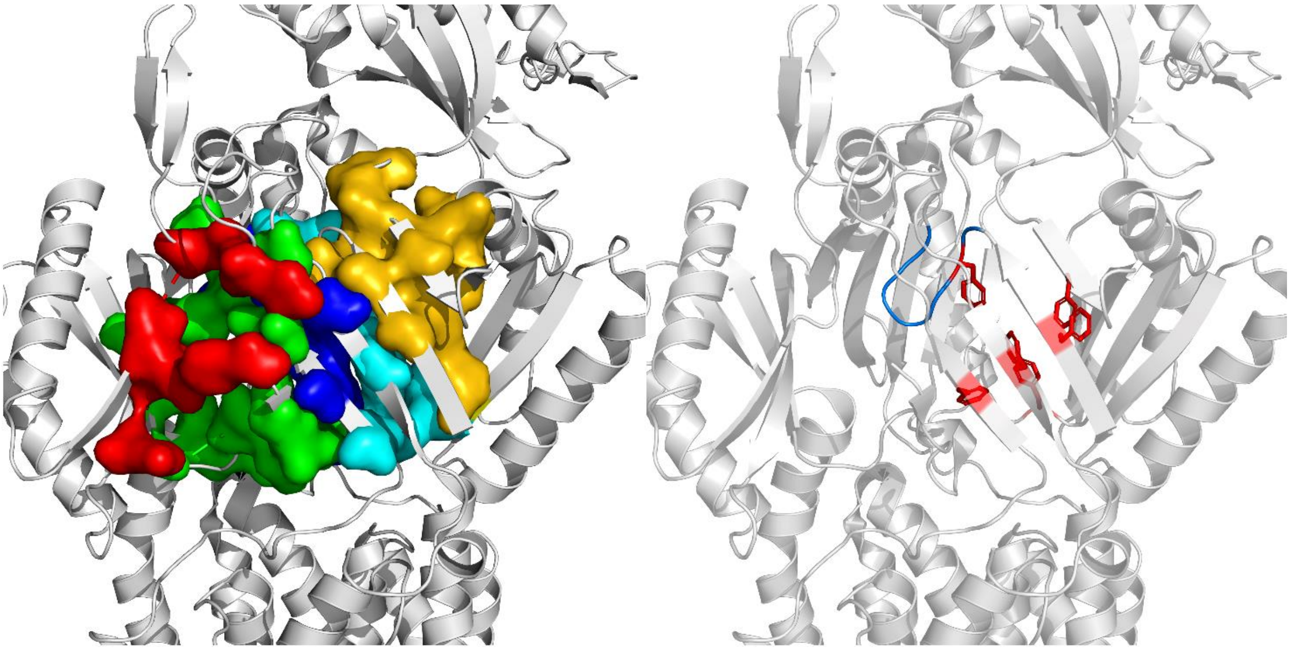

**Structural analysis of docking complexes for representative Rempex compounds.**

To identify differences in key residues and binding modes of the Rempex classes, and thus select good starting configurations for MD simulations, we analyzed in detail the MexB/Rempex complexes representative of each sub-region. **Figures S1** and **S2** compare the predicted binding modes of SUB58, EPI18, EPI-S32 and AVD108 in the $AP_L$ and $DP_T$, respectively. **Figure S1A** shows the representative docking pose of SUB58, with the 3-fluoro-4-(trifluoromethyl)benzyl group that lies on the hydrophobic part of the $AP_L$, in a region delimited by L564, P669, V671, L674 and L861, while the polycationic moiety faces the periplasm. The polyamide linker allows the 2-azaniumyl-ethyl branches to engage stabilizing hydrogen bonds, anchoring the SUB58 to the pocket entrance. **Figure S1B** shows the inhibitor EPI18 assuming a configuration almost superimposable with that of SUB58 in terms of general orientation of the *p*-trifluoromethylphenyl (inward) and the 2-azaniumyl-ethyl groups (outward). The amide substituent at the quinoline ring of EPI-S32 forms a hydrogen bond with E829, which seems to contribute to the stabilization of the compound at the $AP_L$ (**Figure S1C**). AVD108 presents an interaction pattern like that of the other compounds (with the polar amines pointing towards the periplasm and the aromatic rings facing the inner portion of the pocket) (**Figure S1D**).

**FIGURE S2.** Representative modes of binding at the entrance channel of the L monomer of MexB. (**A**) SUB58 (blue, score: -10.6 kcal/mol). (**B**) EPI18 (orange, score: -11.9 kcal/mol). (**C**) EPI-S32 (gray, score: -11.6 kcal/mol). (**D**) AVD108 (yellow, score: -12.9 kcal/mol). Interactions with residues are highlighted as dotted lines.

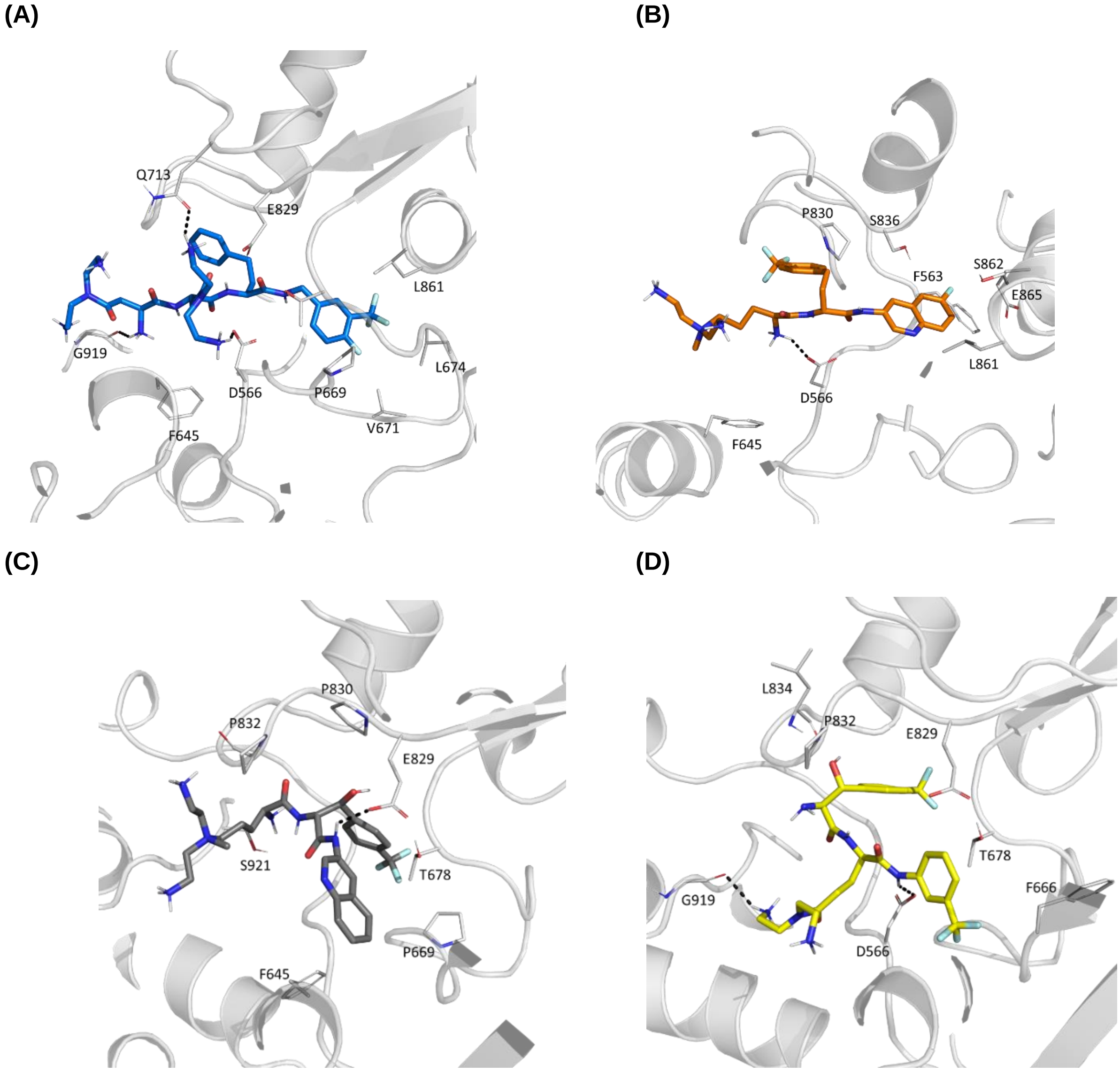


**Figure S2** compares the representative binding modes of SUB58 (**S2A**), EPI18 (**S2B**), and EPI-S32 (**S2C**) in the $DP_T$ (AVD108 was not simulated at the $DP_T$). SUB58 elicits both hydrophobic (F610, F617) and polar (Q46, N718, E825) contacts. Although the 3-fluoro-4-(trifluoromethyl)benzyl group of SUB58 lies in the hydrophobic trap, no *π-π* interactions were found. EPI18 is mainly involved in hydrophobic interactions (F136, F573, F615, F617 belonging to or near the hydrophobic trap), and in one hydrogen bond with T91. Interestingly, EPI-S32 establishes different interactions in the $DP_T$. The specific chirality constrains the compound in terms of interaction with the hydrophobic trap, where the quinoline and the *p*-trifluoromethylphenyl contact several phenylalanine residues, but only the former establishes clear π-π stacking with F628 and F615, while the latter point towards V139, P326 and M630. A hydrogen bond is present, involving the cationic group and F617.

**FIGURE S3.** Same as Figure 2 for the representative binding modes at the T monomer of MexB. (**A**) SUB58 (blue, score -13.6 kcal/mol). (**B**) EPI18 (orange, score -13.7 kcal/mol). (**C**) EPI-S32 (gray, score: -13.8 kcal/mol). Interactions with residues are highlighted as dotted lines.

**(A)**

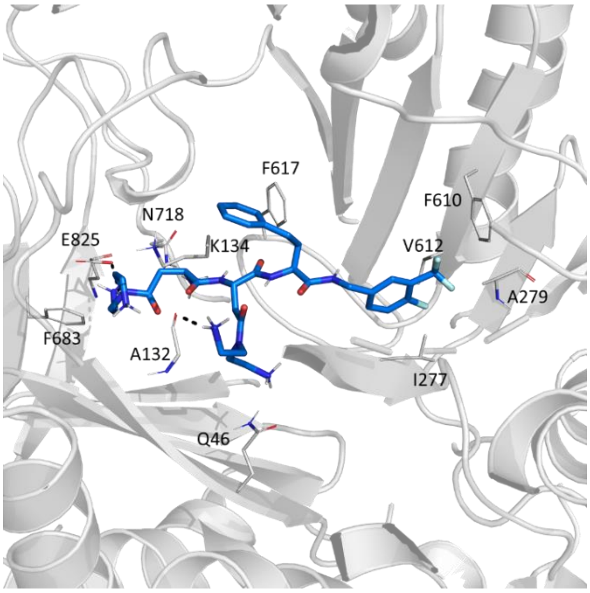


**(B)**

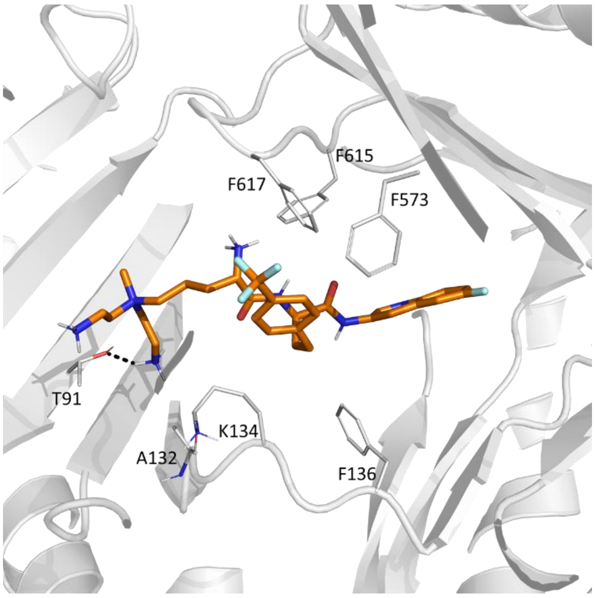


**(C)**

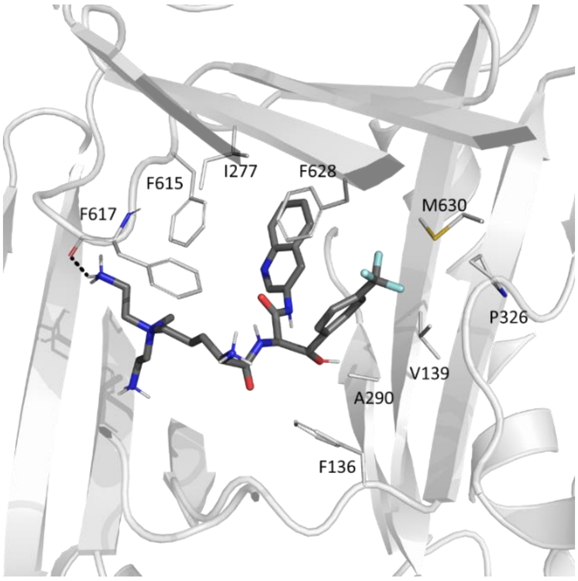

**FIGURE S4.** Mean RMSD values (Å) with the corresponding standard deviations of the ten MD replicas, performed at the $AP_L$ (left) and $DP_T$ (right). (**A**) SUB58, (**B**) EPI18, (**C**) EPI-S32 and (**D**) AVD108.

$AP_L$ $DP_T$

(**A**) SUB58

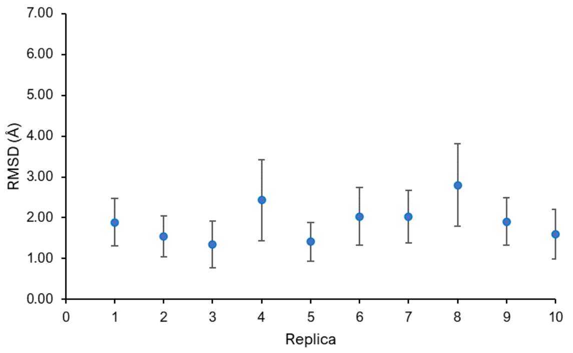


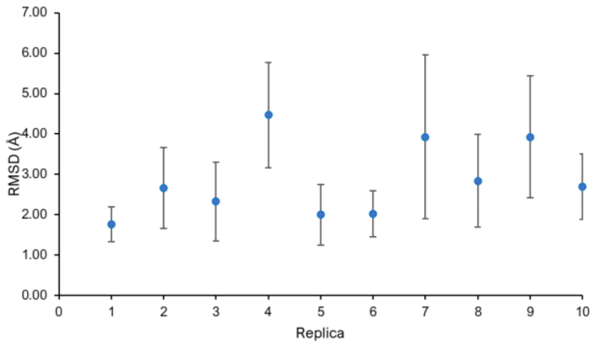


(**B**) EPI18

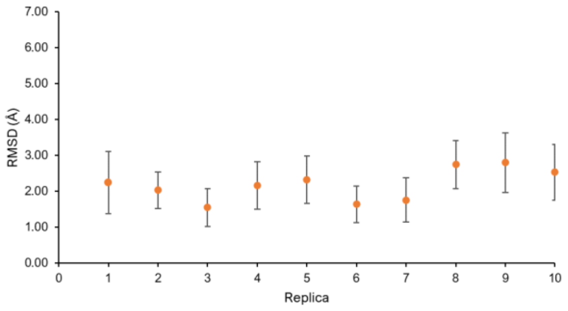


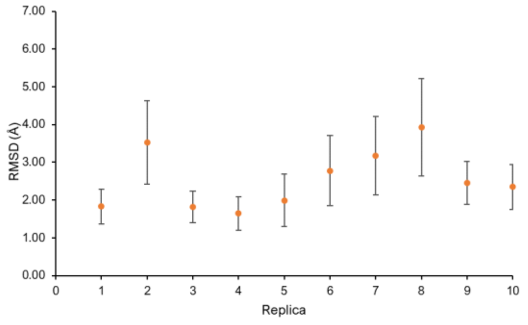


(**C**) EPI-S32

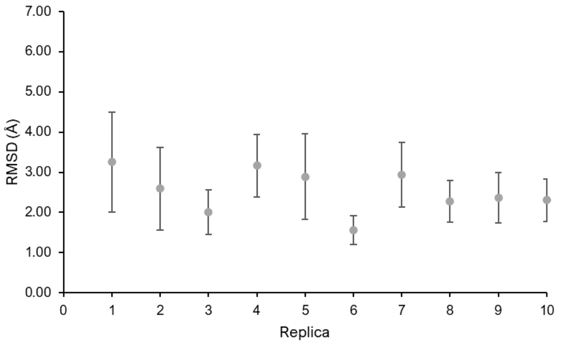


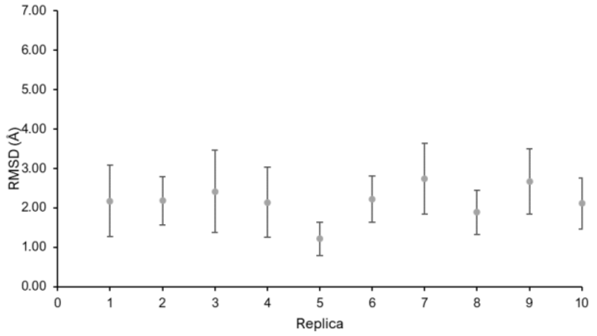


(**D**) AVD108

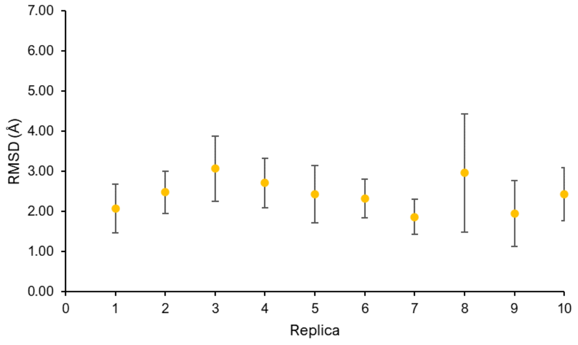

**FIGURE S5.** Pharmacophore groups of Rempex compounds used in contact H-bond water-mediated bridge analyses. (**A**) SUB58, (**B**) EPI18, (**C**) EPI-S32, and (**D**) AVD108. Spheres are colored according to the type of pharmacophore assigned to the functional group: purple for aromatic, cyan for H-donor, blue for H-donor and positively charged, orange for H-acceptor, and green for halogen.

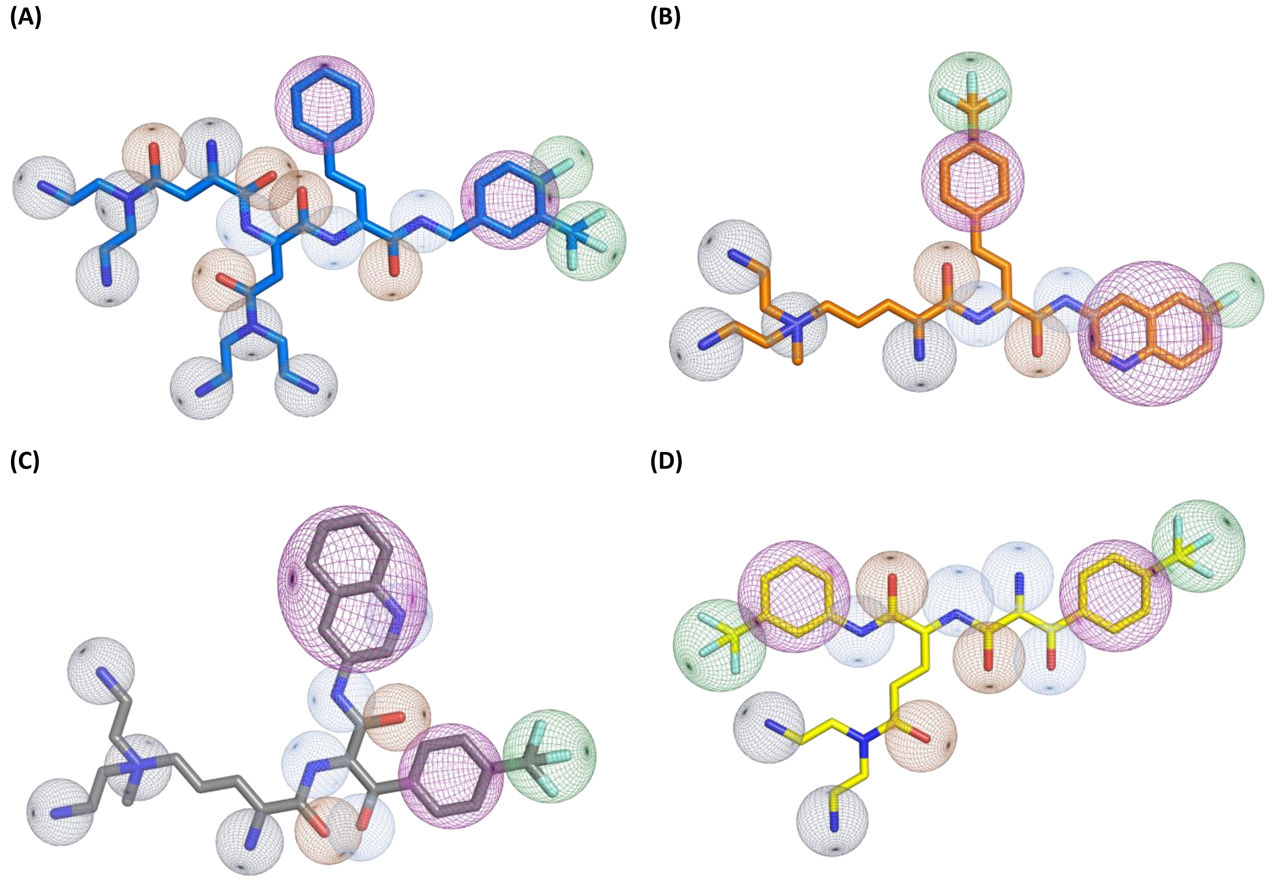


**Figure S6**. Relative levels of expression of MexB variants. Membrane fractions were isolated by ultracentrifugation and proteins were resolved using 10% SDS-PAGE. MexB variants were visualized by immunoblotting with monoclonal anti-His tag antibody (Sigma) and the intensity of the bands was measured by densitometry. The expression of MexB variants is shown as a percent of the expression of MexB WT loaded and analyzed on the same immunoblots. Error bars are SE (n=2).

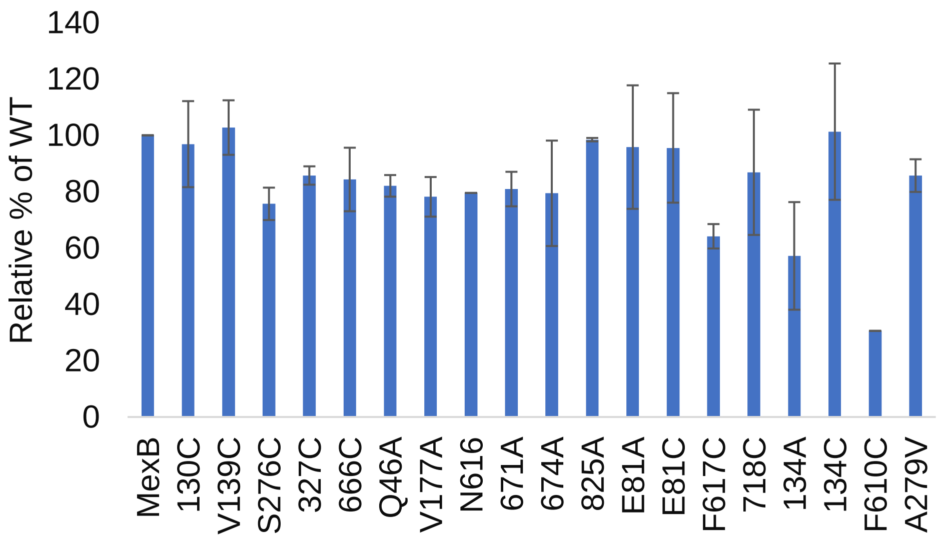

**Figure S7.** Growth inhibition of PA2859(Pore) cells producing the indicated MexB variants by EPI-S32. Cells carrying MexB and E81A were grown in LB broth supplemented with 32 mg/L of NOV and increasing concentrations of EPI-S32. Cells carrying E81C and V671A were grown at 16 mg/L of NOV and increasing concentrations of EPI-S32. Optical density was measured after 24 hrs of incubation and plotted as a function of EPI-S32 concentration. Error bars SD (n=2).

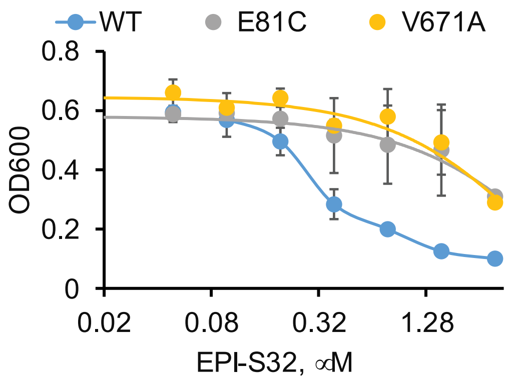


**Table S1. MexB residues of the five ligand binding regions.**

| DP Cave | DP Groove | Interface | AP Outer | AP Inner |
|---|---|---|---|---|
| GLN46 | VAL177 | SER79 | GLN577 | GLN575 |
| THR89 | PHE178 | GLU81 | MET662 | PHE666 |
| ARG128 | SER180 | THR91 | PHE664 | ALA667 |
| THR130 | GLN273 | LYS134 | ARG716 | PRO668 |
| ASN135 | ASP274 | GLY179 | | VAL671 |
| PHE136 | SER276 | ALA279 | | LEU674 |
| VAL139 | ILE277 | PHE573 | | ASN676 |
| GLN176 | PHE610 | ASN616 | | ASP681 |
| LYS292 | VAL612 | PHE617 | | ASN718 |
| TYR327 | PHE615 | | | GLU825 |
| ARG620 | | | | |
| PHE628 | | | | |

**Table S2.** Persistence of direct interaction (%) between Rempex compounds and MexB residues during MD simulations, at both the $AP_L$ and $DP_T$.

| | $AP_L$ persistence of interaction (%) | | | |
|---|---|---|---|---|
| **Resid** | **SUB58** | **AVD108** | **EPI-S32** | **EPI18** |
| I559 | 0.0 | 1.3 | 0.0 | 0.0 |
| P560 | 1.5 | 0.0 | 0.3 | 0.6 |
| T561 | 0.0 | 0.0 | 0.9 | 0.3 |
| A562 | 4.0 | 3.6 | 3.1 | 4.1 |
| F563 | 2.5 | 2.1 | 1.2 | 5.9 |
| L564 | 4.6 | 0.8 | 0.0 | 2.9 |
| P565 | 4.2 | 13.3 | 0.0 | 1.2 |
| D566 | 23.5 | 7.8 | 18.2 | 11.8 |
| E567 | 9.3 | 0.0 | 0.0 | 0.0 |
| D568 | 0.4 | 0.0 | 0.0 | 0.0 |
| F645 | 0.4 | 7.6 | 13.2 | 6.2 |
| E646 | 0.0 | 0.8 | 0.0 | 0.0 |
| A648 | 0.0 | 1.9 | 3.1 | 0.0 |
| K649 | 0.0 | 1.1 | 0.0 | 0.0 |
| Q652 | 0.0 | 1.1 | 0.0 | 0.0 |
| F664 | 0.0 | 0.0 | 2.8 | 0.0 |
| A665 | 0.0 | 0.0 | 2.8 | 0.0 |
| F666 | 0.0 | 1.7 | 3.7 | 0.0 |
| A667 | 0.0 | 3.8 | 6.8 | 0.9 |
| P668 | 0.0 | 1.9 | 3.4 | 0.0 |
| P669 | 0.0 | 1.9 | 3.7 | 3.2 |
| L674 | 0.0 | 0.0 | 0.0 | 2.6 |
| T678 | 3.0 | 10.0 | 4.9 | 4.4 |
| G679 | 2.7 | 1.7 | 0.0 | 1.8 |
| F680 | 4.7 | 0.0 | 0.0 | 2.1 |
| R714 | 1.5 | 0.0 | 3.7 | 0.0 |
| E829 | 7.6 | 0.0 | 2.8 | 14.4 |
| P830 | 6.6 | 5.3 | 3.1 | 8.8 |
| A831 | 0.8 | 1.7 | 2.5 | 0.9 |
| P832 | 0.8 | 2.3 | 3.4 | 2.6 |
| G833 | 0.0 | 2.1 | 3.4 | 2.9 |
| L834 | 0.0 | 3.8 | 3.1 | 0.0 |
| S835 | 0.0 | 0.0 | 1.2 | 0.0 |
| S836 | 3.8 | 2.7 | 0.0 | 8.8 |
| A839 | 0.0 | 0.2 | 0.0 | 0.0 |
| M840 | 2.1 | 0.9 | 0.0 | 0.0 |
| L861 | 0.0 | 1.7 | 0.0 | 2.9 |
| S862 | 3.8 | 0.0 | 0.0 | 0.0 |
| E865 | 3.4 | 0.0 | 0.0 | 7.4 |
| S868 | 0.8 | 0.0 | 0.0 | 0.0 |
| R918 | 0.0 | 0.2 | 0.0 | 0.0 |
| G919 | 1.9 | 3.8 | 3.1 | 2.9 |
| L920 | 0.6 | 1.9 | 3.1 | 0.0 |
| S921 | 0.4 | 9.8 | 2.8 | 0.3 |
| D923 | 0.0 | 1.3 | 0.0 | 0.0 |
| G996 | 3.8 | 0.0 | 0.0 | 0.0 |
| S997 | 1.5 | 0.0 | 0.0 | 0.0 |

| | $DP_T$ persistence of interaction (%) | | |
|---|---|---|---|
| **Resid** | **SUB58** | **EPI18** | **EPI-S32** |
| A42 | 5.4 | 2.3 | 3.3 |
| I43 | 1.3 | 3.9 | 0.2 |
| A44 | 1.3 | 2.1 | 2.2 |
| V45 | 0.0 | 0.0 | 0.7 |
| Y77 | 0.0 | 4.1 | 0.0 |
| S79 | 3.5 | 1.4 | 2.7 |
| S80 | 0.0 | 0.0 | 2.0 |
| E81 | 9.5 | 13.6 | 9.8 |
| T89 | 1.6 | 4.1 | 0.4 |
| I90 | 0.0 | 0.0 | 0.4 |
| T91 | 11.0 | 5.2 | 2.4 |
| V92 | 1.6 | 2.3 | 0.2 |
| T93 | 1.6 | 4.1 | 0.0 |
| A132 | 1.9 | 4.1 | 3.8 |
| V133 | 2.1 | 0.0 | 0.0 |
| K134 | 5.9 | 0.0 | 5.5 |
| F136 | 4.6 | 0.4 | 0.0 |
| V139 | 2.1 | 0.0 | 0.0 |
| Q176 | 0.0 | 1.7 | 0.0 |
| V177 | 0.0 | 1.9 | 0.0 |
| F178 | 0.3 | 4.1 | 0.0 |
| G179 | 0.0 | 0.6 | 0.0 |
| I277 | 0.0 | 1.7 | 0.0 |
| A290 | 0.0 | 1.4 | 0.0 |
| Y327 | 0.6 | 0.0 | 0.0 |
| F573 | 2.5 | 0.0 | 2.2 |
| F615 | 1.4 | 5.0 | 2.4 |
| N616 | 0.6 | 0.0 | 4.4 |
| F617 | 5.1 | 5.8 | 13.3 |
| A618 | 5.4 | 1.4 | 4.4 |
| G619 | 0.5 | 0.0 | 2.2 |
| R620 | 0.0 | 0.4 | 2.2 |
| M626 | 1.7 | 0.0 | 0.2 |
| F628 | 4.6 | 0.0 | 0.7 |
| F666 | 0.5 | 0.0 | 2.2 |
| P668 | 0.0 | 0.0 | 0.2 |
| L672 | 0.6 | 0.0 | 0.2 |
| L674 | 0.0 | 0.0 | 0.4 |
| G675 | 1.4 | 0.0 | 2.0 |
| N676 | 0.0 | 0.0 | 4.0 |
| F680 | 0.2 | 0.0 | 0.0 |
| D681 | 3.2 | 4.1 | 4.4 |
| L682 | 1.6 | 0.0 | 0.0 |
| F683 | 3.8 | 4.1 | 0.0 |
| N718 | 2.5 | 0.0 | 2.2 |
| K814 | 0.0 | 1.2 | 0.2 |
| E816 | 0.0 | 4.1 | 4.4 |
| R817 | 0.0 | 0.0 | 1.8 |
| E825 | 7.9 | 12.4 | 5.3 |
| L827 | 0.0 | 0.2 | 2.0 |
| T859 | 0.6 | 2.1 | 0.7 |
| G860 | 1.4 | 0.0 | 4.0 |

**Table S3.** Persistence of water-mediated H-bond bridges (%) between Rempex compounds and MexB residues (indirect interaction), at both the $AP_L$ and $DP_T$.

| $AP_L$ persistence of H-bond bridges (%) | | | | |
|---|---|---|---|---|
| **Resid** | **SUB58** | **EPI18** | **EPI-S32** | **AVD108** |
| P560 | 0.01 | 0.22 | 0.00 | 11.13 |
| A562 | 0.03 | 0.00 | 0.00 | 0.00 |
| F563 | 1.30 | 3.72 | 8.71 | 0.13 |
| L564 | 1.51 | 7.02 | 52.26 | 0.19 |
| P565 | 0.00 | 2.28 | 0.85 | 0.02 |
| D566 | 64.09 | 40.33 | 54.12 | 51.00 |
| E567 | 62.00 | 0.80 | 0.16 | 4.47 |
| F645 | 0.09 | 1.20 | 2.80 | 0.88 |
| A648 | 0.00 | 0.02 | 0.00 | 0.00 |
| K649 | 0.09 | 0.79 | 6.15 | 2.59 |
| Q652 | 0.00 | 0.10 | 0.12 | 0.27 |
| A665 | 0.00 | 0.07 | 1.00 | 0.04 |
| A667 | 0.02 | 3.03 | 6.93 | 0.22 |
| P668 | 4.82 | 0.75 | 0.00 | 0.03 |
| L674 | 0.00 | 0.00 | 0.00 | 0.01 |
| N676 | 0.00 | 0.04 | 0.02 | 0.01 |
| T678 | 2.30 | 20.80 | 9.89 | 8.28 |
| G679 | 0.46 | 0.34 | 0.02 | 0.09 |
| F680 | 0.00 | 0.01 | 0.00 | 0.00 |
| R714 | 3.06 | 1.75 | 4.20 | 2.00 |
| E829 | 9.07 | 32.75 | 33.92 | 7.12 |
| P830 | 13.57 | 1.82 | 5.48 | 2.55 |
| A831 | 0.22 | 0.76 | 0.71 | 12.32 |
| P832 | 0.03 | 0.14 | 5.24 | 0.66 |
| G833 | 0.10 | 0.86 | 1.30 | 9.80 |
| L834 | 0.00 | 16.12 | 6.43 | 16.75 |
| S835 | 0.06 | 5.27 | 0.35 | 0.07 |
| S836 | 0.21 | 22.34 | 0.22 | 1.10 |
| L861 | 0.00 | 0.08 | 0.00 | 0.00 |
| S862 | 0.03 | 0.02 | 0.00 | 0.00 |
| E865 | 0.23 | 32.74 | 0.00 | 0.02 |
| G919 | 4.12 | 7.58 | 21.07 | 5.14 |
| L920 | 2.40 | 2.36 | 0.40 | 2.49 |
| S921 | 9.22 | 5.25 | 13.91 | 20.75 |
| D923 | 37.39 | 11.68 | 31.66 | 13.23 |
| G996 | 19.95 | 0.00 | 0.00 | 7.14 |
| S997 | 0.19 | 0.00 | 0.00 | 0.00 |

| $DP_T$ persistence of H-bond bridges (%) | | | |
|---|---|---|---|
| **Resid** | **SUB58** | **EPI18** | **EPI-S32** |
| A42 | 0.85 | 0.00 | 0.71 |
| I43 | 1.27 | 0.00 | 0.00 |
| Q46 | 0.01 | 21.01 | 0.09 |
| Y77 | 0.24 | 1.08 | 1.89 |
| S79 | 2.08 | 3.78 | 5.16 |
| S80 | 0.02 | 1.46 | 4.70 |
| E81 | 42.07 | 57.26 | 16.67 |
| T89 | 1.90 | 12.66 | 4.25 |
| T91 | 6.43 | 2.88 | 7.58 |
| V92 | 8.46 | 0.00 | 0.00 |
| T93 | 5.74 | 0.05 | 0.06 |
| R128 | 0.00 | 0.20 | 0.00 |
| T130 | 0.00 | 0.08 | 0.00 |
| A132 | 2.10 | 0.12 | 11.77 |
| V133 | 12.73 | 0.10 | 5.42 |
| K134 | 12.13 | 1.17 | 10.33 |
| Q176 | 0.53 | 5.16 | 0.05 |
| V177 | 0.00 | 0.11 | 0.03 |
| S276 | 0.00 | 0.09 | 0.00 |
| Y327 | 0.00 | 0.01 | 0.00 |
| N616 | 16.48 | 36.50 | 22.99 |
| F617 | 16.52 | 13.94 | 1.70 |
| A618 | 6.66 | 12.88 | 0.83 |
| G619 | 1.50 | 5.17 | 2.01 |
| R620 | 5.23 | 13.16 | 0.20 |
| L672 | 0.32 | 0.00 | 0.99 |
| G675 | 5.10 | 0.83 | 5.62 |
| N676 | 7.07 | 0.75 | 18.39 |
| D681 | 9.98 | 34.23 | 34.02 |
| L682 | 2.11 | 4.36 | 0.00 |
| F683 | 0.00 | 0.01 | 0.00 |
| N718 | 8.89 | 23.05 | 1.53 |
| K814 | 2.10 | 2.75 | 1.53 |
| E816 | 14.25 | 17.65 | 9.34 |
| R817 | 0.00 | 1.77 | 8.17 |
| E825 | 51.00 | 31.15 | 8.54 |
| T859 | 14.92 | 11.75 | 0.52 |
| G860 | 1.06 | 0.26 | 0.12 |

**Table S4.** MIC values (mg/liter ) of the four representative Rempex compounds.

| Rempex | SUB58 | EPI18 | EPI-S32 | AVD108 |
|---|---|---|---|---|
| PAO1[a] | 200 | 6.25 | 100 | 6.25 |
| PAO1-Pore[a] | 100 | 6.25 | 50 | 6.25 |
| PΔ6[a] | 25 | 3.12 | 3.12 | 6.25 |
| PΔ6-Pore[a] | 1.56 | 3.12 | 3.12 | 6.25 |
| PAO2859(Pore, pMexB) | 100 | 12.5 | 200 | ND |
| PAO2859(Pore, pUCP22) | 25 | 12.5 | 25 | ND |

[a], Data were retrieved from Mehla *et al.*[1]
